# Estimating Transfer Entropy in Continuous Time Between Neural Spike Trains or Other Event-Based Data

**DOI:** 10.1101/2020.06.16.154377

**Authors:** David P. Shorten, Richard E. Spinney, Joseph T. Lizier

## Abstract

Transfer entropy (TE) is a widely used measure of directed information flows in a number of domains including neuroscience. Many real-world time series in which we are interested in information flows come in the form of (near) instantaneous events occurring over time, including the spiking of biological neurons, trades on stock markets and posts to social media. However, there exist severe limitations to the current approach to TE estimation on such event-based data via discretising the time series into time bins: it is not consistent, has high bias, converges slowly and cannot simultaneously capture relationships that occur with very fine time precision as well as those that occur over long time intervals. Building on recent work which derived a theoretical framework for TE in continuous time, we present an estimation framework for TE on event-based data and develop a *k*-nearest-neighbours estimator within this framework. This estimator is provably consistent, has favourable bias properties and converges orders of magnitude more quickly than the discrete-time estimator on synthetic examples. We also develop a local permutation scheme for generating null surrogate time series to test for the statistical significance of the TE and, as such, test for the conditional independence between the history of one point process and the updates of another — signifying the lack of a causal connection under certain weak assumptions. Our approach is capable of detecting conditional independence or otherwise even in the presence of strong pairwise time-directed correlations. The power of this approach is further demonstrated on the inference of the connectivity of biophysical models of a spiking neural circuit inspired by the pyloric circuit of the crustacean stomatogastric ganglion, succeeding where previous related estimators have failed.

**AUTHOR SUMMARY:** Transfer Entropy (TE) is an information-theoretic measure commonly used in neuroscience to measure the directed statistical dependence between a source and a target time series, possibly also conditioned on other processes. Along with measuring information flows, it is used for the inference of directed functional and effective networks from time series data. The currently-used technique for estimating TE on neural spike trains first time-discretises the data and then applies a straightforward or “plug-in” information-theoretic estimation procedure. This approach has numerous drawbacks: it is very biased, it cannot capture relationships occurring on both fine and large timescales simultaneously, converges very slowly as more data is obtained, and indeed does not even converge to the correct value. We present a new estimator for TE which operates in continuous time, demonstrating via application to synthetic examples that it addresses these problems, and can reliably differentiate statistically significant flows from (conditionally) independent spike trains. Further, we also apply it to more biologically-realistic spike trains obtained from a biophysical model of the pyloric circuit of the crustacean stomatogastric ganglion; our correct inference of the underlying connection structure here provides an important validation for our approach where similar methods have previously failed

## I. INTRODUCTION

In analysing time series data from complex dynamical systems, such as in neuroscience, it is often useful to have a notion of information flow. We intuitively describe the activities of brains in terms of such information flows: for instance, information from the visual world must flow to the visual cortex where it will be encoded [1]. Conversely, information coded in the motor cortex must flow to muscles where it will be enacted [2].

Transfer entropy (TE) [3, 4] has become a widely accepted measure of such flows. It is defined as the mutual information between the past of a source time-series process and the present state of a target process, conditioned on the past of the target. More specifically (in discrete time), the transfer entropy rate [5] is:

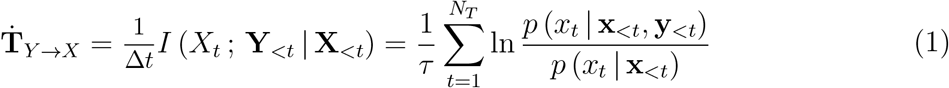

Here the information flow is being measured from a source process *Y* to a target *X*, *I*(· ; · | ·) is the conditional mutual information [6], *x_t_* is the current state of the target, **x**_<*t*_ is the history of the target, **y**_<*t*_ is the history of the source, Δ*t* is the interval between time samples (in units of time), *τ* is the length of the time series and *N_T_* = *τ/*Δ*t* is the number of time samples. The histories **x**_<*t*_ and **y**_<*t*_ are usually captured via embedding vectors, e.g. **x**_<*t*_ = **x**_*t*−*m*:*t*−1_ = {*x_t_*_−*m*_, *x*_*t−m*+1_, … , *x*_*t−1*_}. Recent work [5] has highlighted the importance of normalising the TE by the width of the time bins, as above, such that it becomes a *rate*, in order to ensure convergence in the limit of small time bin size.

It is also possible to condition the TE on additional processes [4]. Given additional processes 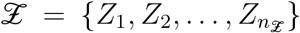 with histories 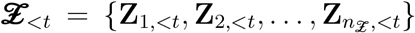, we can write the conditional TE rate as

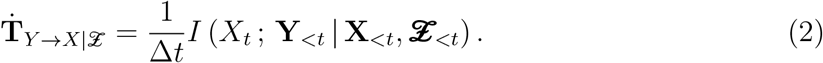

When combined with a suitable statistical significance test, the conditional TE can be used to show that the present state of *X* is conditionally independent of the past of *Y* – conditional on the past of *X* and of the conditional processes 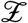. Such conditional independence implies the absence of an active causal connection between *X* and *Y* under certain weak assumptions [7, 8], and TE is widely used for inferring directed functional and effective network models [9–12] (and see [4, Sec. 7.2]).

TE has received widespread application in neuroscience [13, 14]. Uses have included the network inference as above, as well as measuring the direction and magnitude of information flows [15, 16] and determining transmission delays [17]. Such applications have been performed using data from diverse sources such as MEG [18, 19], EEG [20], fMRI [21], electrode arrays [22], calcium imaging [11] and simulations [23].

Previous applications of TE to spike trains [22, 24–34] and other types of event-based data [35], including for the purpose of network inference [11, 36, 37], have relied on time discretisation. As shown in Fig. 1, the time series is divided into small bins of width Δ*t*. The value of a sample for each bin could then be assigned as either binary - denoting the presence or absence of events (spikes) in the bin - or a natural number denoting the number of events (spikes) that fell within the bin. A choice is made as to the number of time bins, *l* and *m*, to include in the source and target history embeddings **y**_<*t*_ and **x**_<*t*_. This results in a finite number of possible history embeddings. For a given combination **x**_<*t*_ and **y**_<*t*_, the probability of the target’s value in the current bin conditioned on these histories, *p* (*x*_*t*_ | **x**_<*t*_, **y**_<*t*_), can be directly estimated using the so-called plugin (histogram) estimator. The probability of the target’s value in the current bin conditioned on only the target history, *p* (*x_t_* | **x**_<*t*_), can be estimated in the same fashion. From these estimates the TE can be calculated in a straightforward manner via (1). (See Results for a description of the application of the discrete time TE estimator to synthetic examples including spiking events from simulations of model neurons.)

**FIG. 1:**
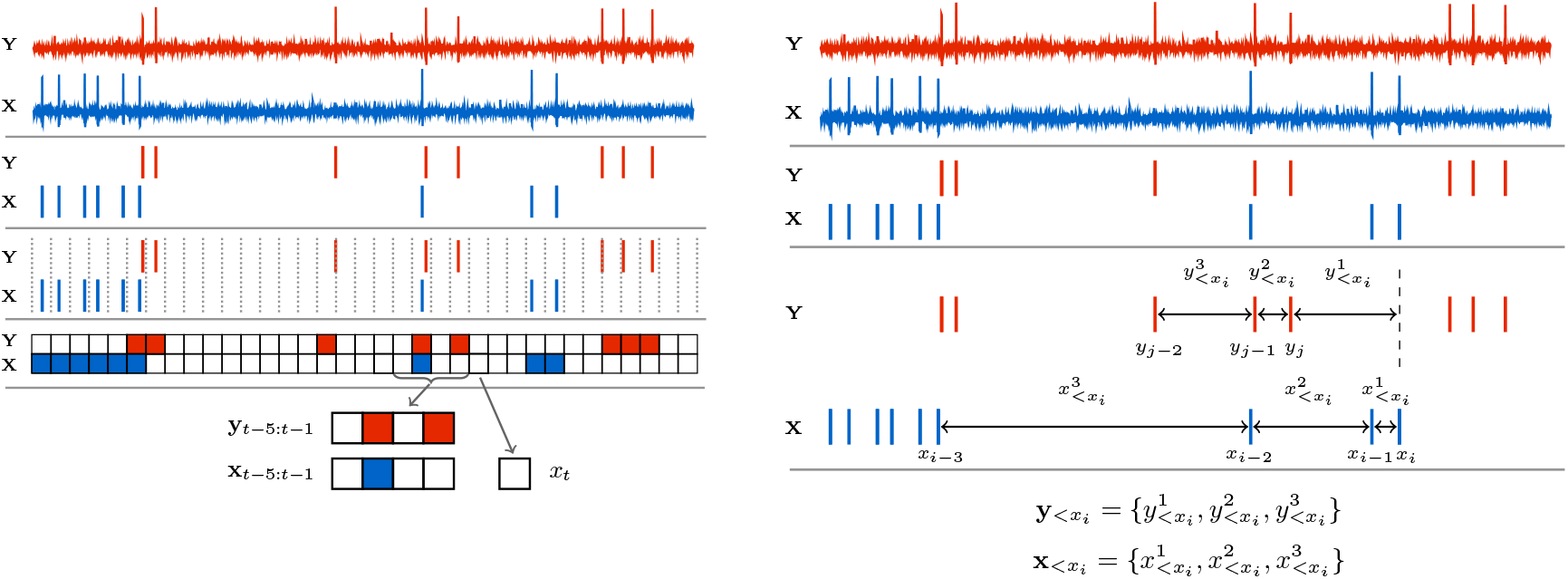
Diagrams highlighting the differences in the embeddings used by the discrete and continuous-time estimators. The discrete-time estimator (left) divides the time series into time bins. A binary value is assigned to each bin denoting the presence or absence of a (spiking) event – alternatively, this could be a natural number to represent the occurrence of multiple events. The process is thus recast as a sequence of binary values and the history embeddings (**x**_*t*−5:*t*−1_ and **y**_*t*−5:*t*−1_) for each point are binary vectors. The probability of an event occurring in a bin, conditioned on its associated history embeddings, is estimated via the so-called plugin (histogram) estimator. Conversely, the continuous-time estimator (right) performs no time binning. History embeddings 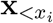 and 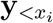 for events or 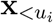 and 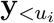 for arbitrary points in time (not shown, see Methods) are constructed from the raw interspike intervals. This approach estimates the TE by comparing the probabilities of the history embeddings of the target processes’ history as well as the joint history of the target and source processes at both the (spiking) events and arbitrary points in time.

There are two large disadvantages to this approach [5]. If the process is genuinely occurring in discrete time, then the estimation procedure just described is consistent. That is, it is guaranteed to converge to the true value of the TE in the limit of infinite data. However, if we are considering a fundamentally continuous-time process, such as a neuron’s action potential, then the lossy transformation of time discretisation will result in an inaccurate estimate of the TE. Thus, in these cases, any estimator based on time discretisation is not consistent. Secondly, whilst the loss of resolution of the discretization will decrease with decreasing bin size Δ*t*, this requires larger dimensionality of the history embeddings to capture correlations over similar time intervals. This increase in dimension will result in an exponential increase in the state space size being sampled and therefore the data requirements. This renders the estimation problem intractable for the typical dataset sizes present in neuroscience.

In practice then, the application of transfer entropy to event-based data such as spike trains has required a trade-off between fully resolving interactions that occur with fine time precision and capturing correlations that occur across long time intervals. There is substantial evidence that spike correlations at the millisecond and sub-millisecond scale play a role in encoding visual stimuli [38, 39], motor control [40] and speech [41]. On the other hand, correlations in spike trains exist over lengths of hundreds of milliseconds [42]. A discrete-time TE estimator cannot capture both of these classes of effects simultaneously.

Recent work by Spinney et al. [5] derived a continuous-time formalism for TE. It was demonstrated that, for stationary point processes such as spike trains, the pairwise TE rate is given by:

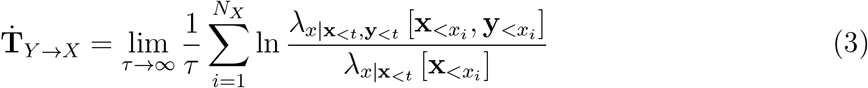

It is worth emphasizing that, in this context, the processes *X* and *Y* are series of the time points *x_i_* of events *i*. This is contrasted with (1), where *X* and *Y* are series of values at the sampled time points *t_i_*. To avoid confusion we use the notation that the *y_i_* ∈ *Y* are the raw time points and 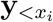 is some representation of the history of *Y* observed at the time point *x_i_* (see Methods). Here, 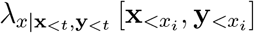 is the instantaneous firing rate of the target conditioned on the histories of the target 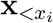 and source 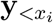 at the time points *x_i_* of the events in the target process. 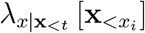 is the instantaneous firing rate of the of the target conditioned on its history alone, ignoring the history of the source. Note that 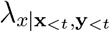 and 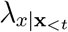 are defined at all points in time and not only at target events. Equation (3) can easily be adapted to the conditional case:

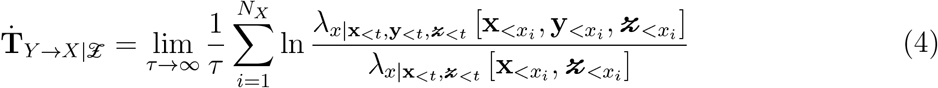

Here, 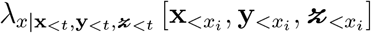 is the instantaneous firing rate of the target conditioned on the histories of the target 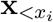, source 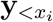 and other possible conditioning processes 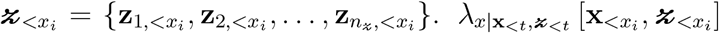 is the instantaneous firing rate of the of the target conditioned on the histories of the target and the additional conditioning processes, ignoring the history of the source.

Crucially, it was demonstrated by Spinney et al., and later shown more rigorously by Cooper and Edgar [43], that if the discrete-time formalism of the TE (in (1)) could be properly estimated as lim_*δt*→0_, then it would converge to the same value as the continuous-time formalism. This is due to the contributions to the TE from the times between target events cancelling. Yet there are two important distinctions in the continuous-time formalism which hold promise to address the consistency issues of the discrete-time formalism. First, the basis in continuous time allows us to represent the history embeddings by inter-event intervals, suggesting the possibility of jointly capturing subtleties in both short and long time-scale effects that has evaded discrete-time approaches. (See Fig. 1 for a diagrammatic representation of these history embeddings, contrasted with the traditional way of constructing histories for the discrete-time estimator). Second, note the important distinction that the sums in (3) and (4) are taken over the *N_X_* (spiking) events in the target during the time-series over interval *τ*; this contrasts to a sum over all time-steps in the discrete-time formalism. An estimation strategy based on (3) and (4) would only be required to calculate quantities at events, ignoring the ‘empty’ space in between. This implies a potential computational advantage, as well as eliminating one source of estimation variability.

These factors all point to the advantages of estimating TE for event-based data using the continuous-time formalism in (3) and (4). This paper presents an empirical approach to doing so. The estimator (presented in Methods) operates by considering the probability densities of the history embeddings observed at events and contrasts these with the probability densities of those embeddings being observed at other (randomly sampled) points. This approach is distinct in utilising a novel Bayesian inversion on (4) in order to operate on these probability densities of the history embeddings, rather than making a more difficult direct estimation of spike rates. This further allows us to utilise *k*-Nearest-Neighbour (*k*NN) estimators for entropy terms based these probability densities, bringing known advantages of consistency, data efficiency, low sensitivity to parameters and known bias corrections. By combining these entropy estimators, and adding a further bias-reduction step, we arrive at our proposed estimator. The resulting estimator is consistent (see Methods section IV A 5) and is demonstrated on synthetic examples in Results to be substantially superior to estimators based on time discretisation across a number of metrics.

These surrogate samples have historically been created by either shuffling the original source samples or by shifting the source process in time. However, this results in surrogates which conform to an incorrect null hypothesis that the transitions in the target are completely independent of the source histories. That is, they conform to the factorisation 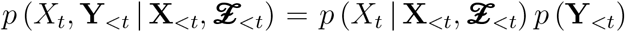. This leads to the possibility that a process could possess correlations that deviate from such a null which are not measured by TE.

As shown in Results, this can lead to incredibly high false positive rates in certain settings such as the presence of strong common driver effects. Therefore, in order to have a suitable significance test for use in conjunction with the proposed estimator, we also present (in Methods section IV B) an adaptation of a recently proposed local permutation method [8] to our specific case. This adapted scheme produces surrogates which conform to the correct null hypothesis of conditional independence of the present of the target and the source history, given the histories of the target and further conditioning processes. This is the condition that 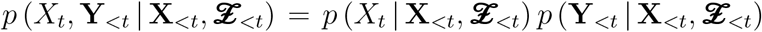. All deviations from such a condition are contributions to the transfer entropy and so it is deemed to be the appropriate null.

We show in Results that the combination of the proposed estimator and surrogate generation method is capable of correctly distinguishing between zero and non-zero information flow in cases where the history of the source might have a strong pairwise correlation with the occurrence of events in the target, but is nevertheless conditionally independent. The combination of the discrete-time estimator and a traditional method of surrogate generation is shown to be incapable of making this distinction. Further, we demonstrate that the proposed combination is capable of inferring the connectivity of a simple circuit of bio-physical model neurons inspired by the crustacean stomatogastric ganglion, a task which the discrete-time estimator was shown to be incapable of performing. This provides strong impetus for the application of the proposed techniques to investigate information flows in spike-train data recorded from biological neurons.

## II. RESULTS

The first two subsections here present the results of the continuous-time estimator applied to two different synthetic examples for which the ground truth value of the TE is known. The first example considers independent processes where 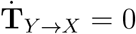, whilst the second examines coupled Poisson processes with a known, non-zero 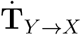. The continuous-time estimator’s performance is also contrasted with that of the discrete-time estimator. The emphasis of these sections is on properties of the estimators: their bias, variance and consistency (see Methods).

The third and fourth subsections present the results of the combination of the continuous-time estimator and the local permutation surrogate generation scheme applied to two examples: one synthetic and the other a biologically plausible model of neural activity. The comparison of the estimates to a population of surrogates produces *p*-values for the statistical significance of the TE values being distinct from zero and implying a directed relationship. The results are compared to the known connectivity of the studied systems. These *p*-values could be translated into other metrics such as ROC curves and false-positive rates, but we choose to instead visualise the distributions of the *p*-values themselves. The combination of the discrete-time estimator along with a traditional method for surrogate generation (time shifts) is also applied to these examples for comparison.

### A. No TE Between Independent Homogeneous Poisson Processes

The simplest example on which we can attempt to validate the estimator is independent homogeneous Poisson processes, where the true value of the TE between such processes is zero.

Pairs of independent homogeneous Poisson processes were generated, each with rate 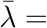 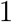, and contiguous sequences of *N_X_* ∈ {1 × 10^2^, 1 × 10^3^, 1 × 10^4^, 1 × 10^5^} target events were selected. For the continuous-time estimator, the parameter *N_U_* for the number of placed sample points was varied (see Methods sections IV A 4 and IV A 5) to check the sensitivity of estimates to this parameter. At each of these numbers of target events *N_X_*, the averages are taken across 1000, 100, 20 and 20 tested process pairs respectively.

Fig. 2 shows the results of these runs for the continuous-time estimator, using various parameter settings. In all cases, the Manhattan (*ℓ*_1_) norm is used as the distance metric and the embedding lengths are set to *l_X_* = *l_Y_* = 1 spike (see Methods sections IV A 4 and IV A 5). For this example, the set of conditioning processes 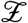 is empty.

**FIG. 2:**
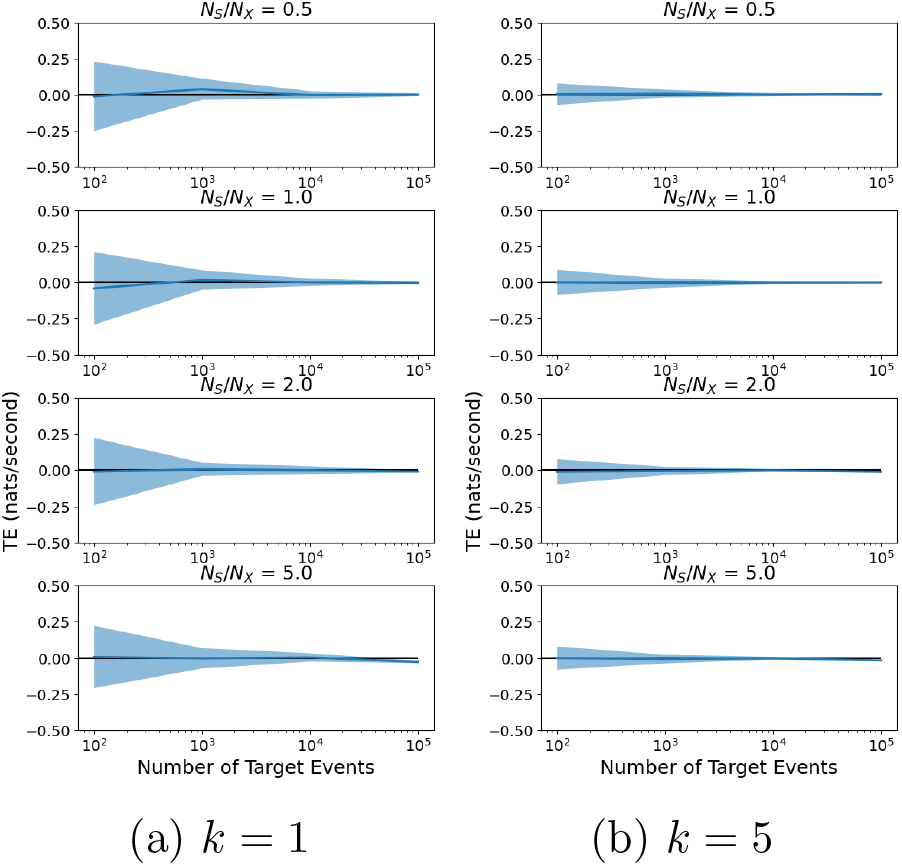
Evaluation of the continuous-time estimator on independent homogeneous Poisson processes. The blue line shows the average TE rate across multiple runs and the shaded area shows the standard deviation. Plots are shown for two different values of *k* nearest neighbours, and four different values of the ratio of the number of sample points to the number of events *N_U_ /N_X_* (See Methods, sections IV A 4 and IV A 5).

The plots show that the continuous-time estimator converges to the true value of the TE (0). This is a numerical confirmation of its consistency for independent processes. Moreover, it exhibits very low bias (as compared to the discrete-time estimator, Fig. 3) for all values of the *k* nearest neighbours and *N_U_ /N_X_* parameters. The variance is relatively large for *k* = 1, although it is dramatically better for *k* = 5 — this reflects known results for variance of this class of estimators as a function of *k* [44].

**FIG. 3:**
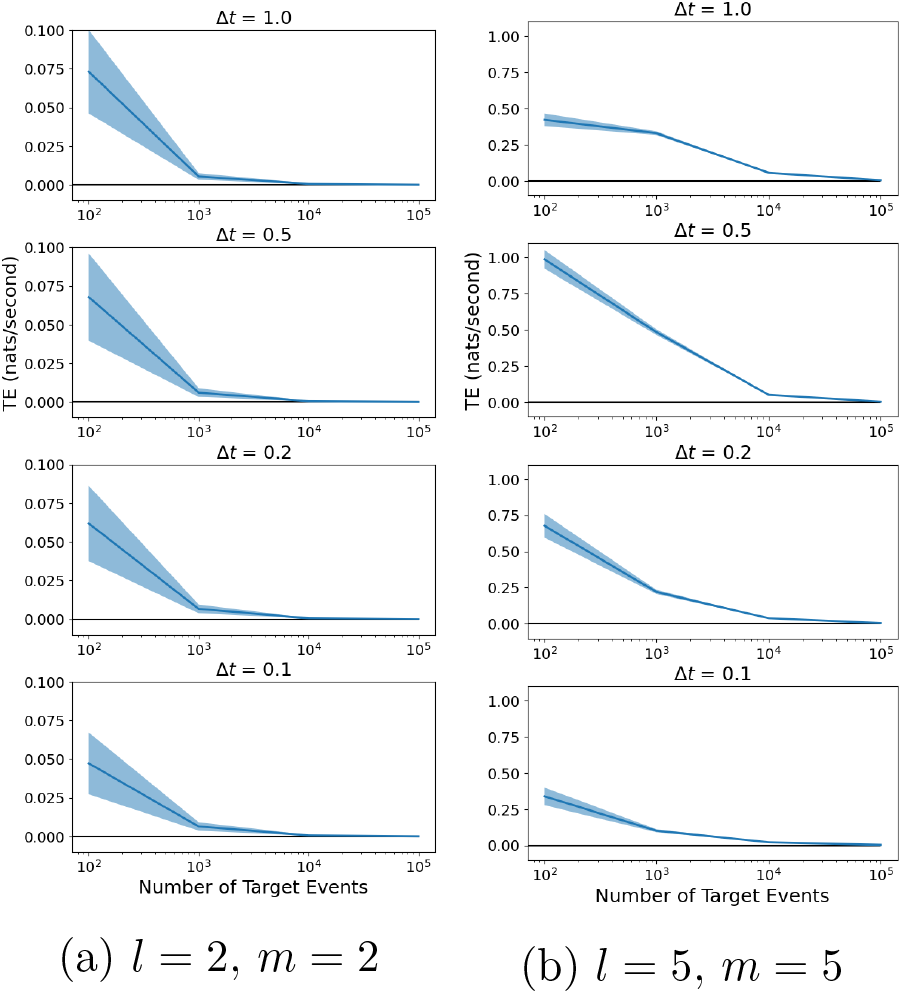
Result of the discrete time estimator applied to independent homogeneous Poisson processes. The solid line shows the average TE rate across multiple runs and the shaded area shows the standard deviation. Plots are shown for four different values of the bin width Δ*t* as well as different source and target embedding lengths, *l* and *m*.

Fig. 3 shows the result of the discrete-time estimator applied to the same independent homogeneous Poisson processes for two different combinations of the source and target history embedding lengths, *l* and *m* time bins, and four different bin sizes Δ*t*. At each of the numbers of target events *N_X_*, the averages are taken across 1000, 100, 100 and 100 tested process pairs respectively. The variance of this estimator on this process is low and comparable to the continuous-time estimator, however the bias is very large and positive for short processes. It would appear that, in the case of these independent processes, it is consistent.

### B. Consistent TE Between Unidirectionally Coupled Processes

The estimators were also tested on an example of unidirectionally coupled spiking processes with a known value of TE (previously presented as example B in [5]). Here, the source process *Y* is an homogoneous Poisson process. The target process *X* is produced as a conditional point process where the instantaneous rate is a function of the time since the most recent source event. More specifically:

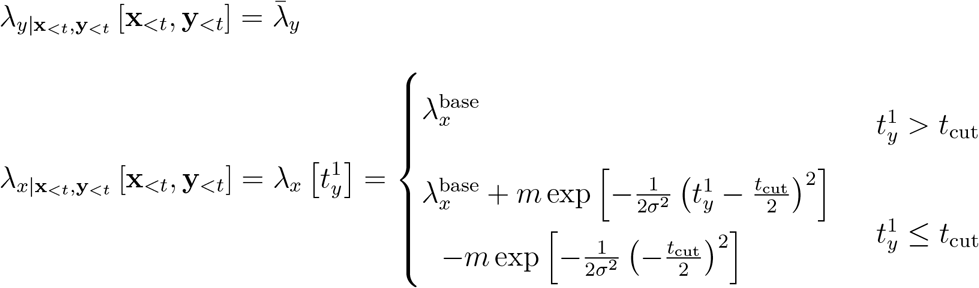

Here, 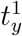 is the time since the most recent source event. As a function of 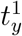, the target spike rate 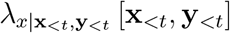 rises from a baseline 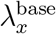 at 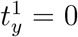 to a peak at 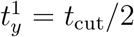, before falling back to the baseline 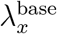 from 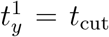 onwards (see Fig. 4a). We simulated this process using the parameter values 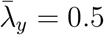, *m* = 5, *t*_cut_ = 1, 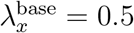 and *σ*^2^ = 0.01. This simulation was performed using a thinning algorithm [45]. Specifically, we first generated the source process at rate 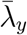. We then generate the target as an homogeneous Poisson process with rate λ_h_ such that 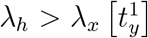 for all values of 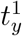. We then went back through all the events in this process and removed each event with probability 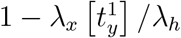. As with the previous example, once a pair of processes had been generated, a contiguous sequence of *N_X_* target events was selected. Tests were conducted for the values of *N_X_* ∈ {1 × 10^2^, 1 × 10^3^, 1 × 10^4^, 1 × 10^5^}. For the continuous-time estimator, the number of placed sample points *N_U_* was set equal to *N_X_* (see Methods sections IV A 4 and IV A 5). At each *N_X_*, the averages are taken over 1000, 100, 20 and 20 tested process pairs respectively. Due to its slow convergence, the discrete-time estimator was also evaluated at a value of *N_X_* = 1 × 10^6^ target spikes, averaged over 20 process pairs.

**FIG. 4:**
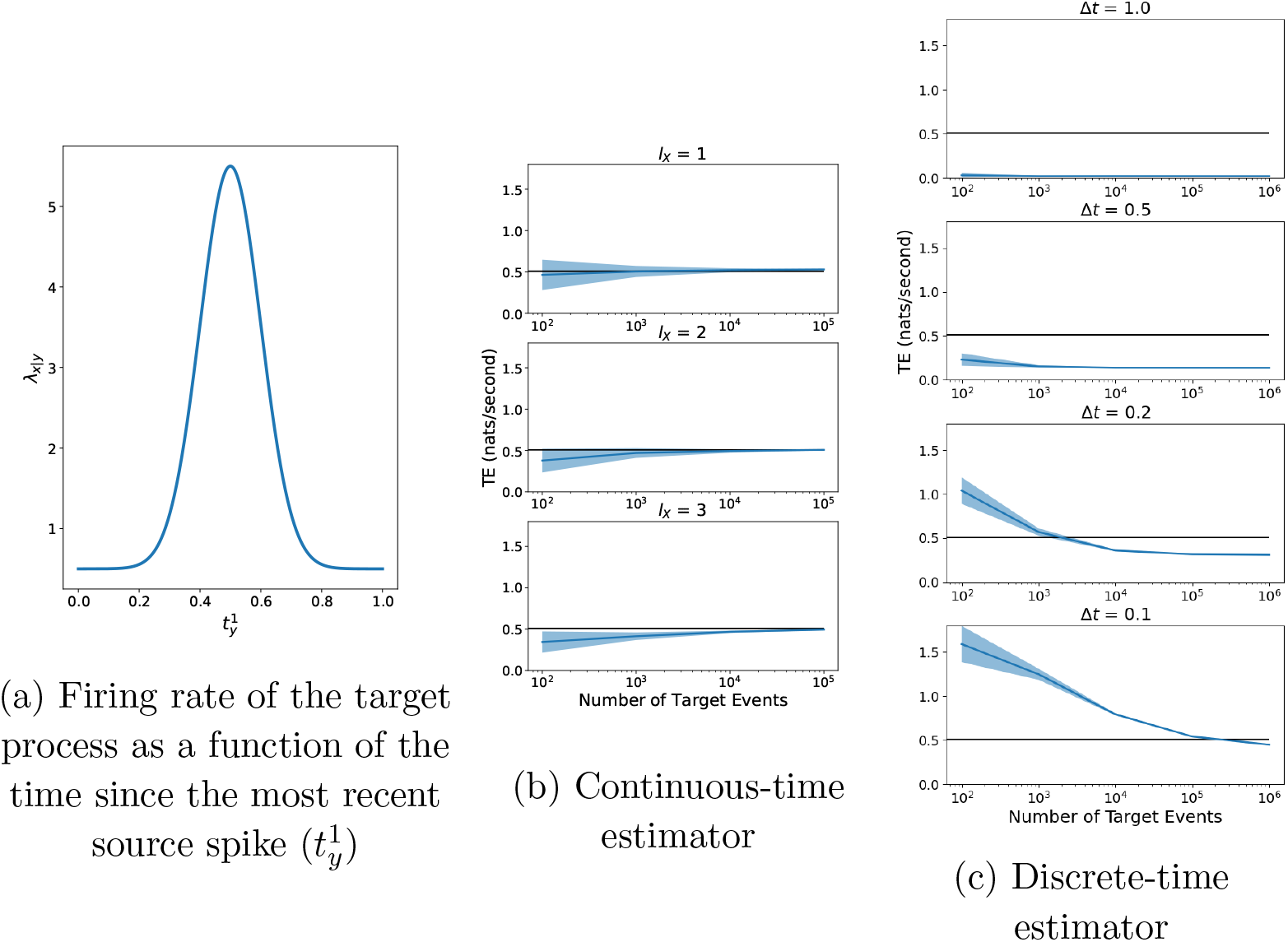
The discrete-time and continuous-time estimators were run on coupled Poisson processes for which the ground-truth value of the TE is known. (a) shows the firing rate of the target process as a function of the history of the source. (b) and (c) show the estimates of the TE provided by the two estimators. The blue line shows the average TE rate across multiple runs and the shaded area shows the standard deviation. The black line shows the true value of the TE. For the continuous-time estimator the parameter values of *N_U_ /N_X_* = 1 and *k* = 4 were used along with the *ℓ*_1_ (Manhattan) norm. Plots are shown for three different values of the length of the target history component *l_X_*. For the discrete-time estimator, plots are shown for four different values of the bin width Δ*t*. The source and target history embedding lengths are chosen such that they extend back one time unit.

Spinney et al. [5] present a numerical method for calculating the TE for this process. For the parameter values used here the true value of the TE is 0.5076 ± 0.001.

Given that we know that the dependence of the target on the source is fully determined by the distance to the most recent event in the source, we used a source embedding length of *l_Y_* = 1. The estimators were run with three different values of the target embedding length *l_X_* ∈ {1, 2, 3} (see Methods sections IV A 4 and IV A 5). For this example, the set of conditioning processes 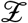 is empty.

Fig. 4b shows the results of the continuous-time estimator applied to the simulated data. We used the value of *k* = 4 and the Manhattan (*ℓ*_1_) norm. The results displayed are as expected in that for a short target history embedding length of *l_X_* = 1 spike, the estimator converges to a slight over-estimate of the TE. The overestimate at shorter target history embedding lengths *l_X_* can be explained in that perfect estimates of the 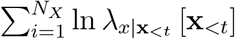 component require full knowledge of the target past within the previous *t*_cut_ = 1 time unit; shorter values of *l_X_* don’t cover this period in many cases, leaving this rate underestimated and therefore the TE overestimated. For longer values of *l_X_* ∈ {2, 3} we see that they converge closely to the true value of the TE. This is a further numerical confirmation of the consistency of the continuous-time estimator.

Fig. 4c shows the results of the discrete-time estimator applied to the same process, run for three different values of the bin width Δ*t* ∈ {1, 0.5, 0.2, 0.1} time units. The number of bins included in the history embeddings was chosen such that they extended one time unit back (the known length of the history dependence). Smaller bin sizes could not be used as this caused the machine on which the experiments were running to run out of memory. The plots are a clear demonstration that the discrete-time estimator is very biased and not consistent. At a bin size of Δ*t* = 0.2 it converges to a value less than half the true TE. Moreover, its convergence is incredibly slow. At the bin size of Δ*t* = 0.1 it would appear to not have converged even after 1 million target events, and indeed it is not even converging to the true value of the TE. The significance of the performance improvement by our estimator is explored further in Discussion.

### C. Identifying Conditional Independence Despite Strong Pairwise Correlations

The existence of a set of conditioning processes under which the present of the target component is conditionally independent of the past of the source implies that, under certain weak assumptions, there is no causal connection from the source to the target [46–48]. More importantly, TE can be used to test for such conditional independence (see Methods), thus motivating its use in network inference. A large challenge faced in testing for conditional independence is correctly identifying “spurious” correlations, whereby conditionally independent components might have a strong pairwise correlation. This problem is particularly pronounced when investigating the spiking activity of biological neurons, populations of which often exhibit highly correlated behaviour through various forms of synchrony [49–51], or common drivers [52, 53]. In this subsection we demonstrate that the combination of the presented estimator and surrogate generation scheme are particularly adept at identifying conditional independence in the face of strong pairwise correlations on a synthetic example. Moreover, the combination of the traditional discrete-time estimator and surrogate generation techniques are demonstrated to be ineffective on this task.

The chosen synthetic example models a common driver effect, where an apparent directed coupling between a source and target is only due to a common parent. In such cases, despite a strong induced correlation between the source history and the occurrence of an event in the target, we expect to measure zero information flow when conditioning on the common driver. Our system here consists of a quasi-periodic ‘mother’ process *M* (the common driver) and two ‘daughter’ processes, *D*_1_ and *D*_2_ (see Fig. 5 for a diagram of the process). The mother process contains events occurring at intervals of *T* + *ξ_M_*, with the daughter processes being noisy copies with each event shifted by an amount 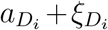. We also choose that 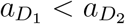; so long as the difference between these 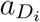 values is large compared to the size of the noise terms, this will ensure that the events in *D*_1_ precede those in *D*_2_. When conditioning on the mother process, the TE from the first daughter to the second, 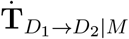, should be 0. However, the history of the source daughter process is strongly correlated with the occurrence of events in the second daughter process — the events in *D*_1_ will precede those in *D*_2_ by the constant amount 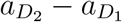 plus a small noise term 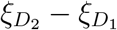.

**FIG. 5:**
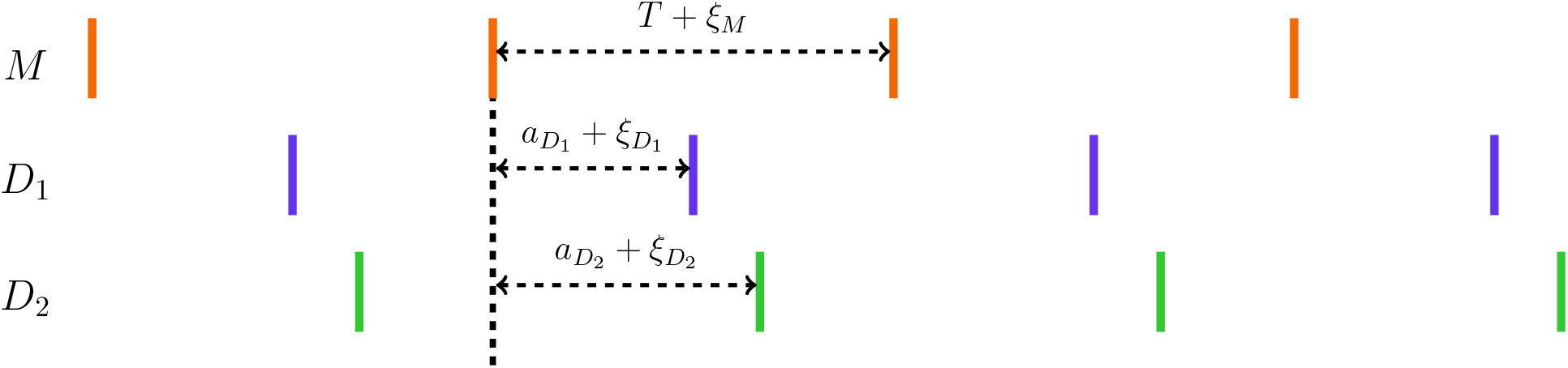
Diagram of the noisy copy process. Events in the mother process *M* occur periodically with intervals *T* + *ξ_M_* (*ξ* is used for noise terms). Events in the daughter processes *D*_1_ and *D*_2_ occur after each event in the mother process, at a distance of 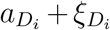

Due to the noise in the system, this level of correlation will gradually break down if we shift the source daughter process relative to the others. This allows us to do two things. Firstly, we can get an idea of the bias of the estimator on conditionally independent processes for different levels of pairwise correlation between the history of the source and events in the target. Second, we can evaluate different schemes of generating surrogate TE distributions as a function of this correlation. We would expect that, for well-generated surrogates which reflect the relationships to the conditional process, the TE estimates on these conditionally independent processes will closely match the surrogate distribution.

We simulated this process using the parameter values of *T* = 1.0, 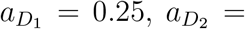 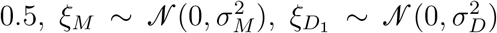 and 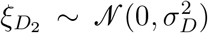, *σ_M_* = 0.05 and *σ_D_* = 0.05. Once the process had been simulated, the source process *D*_1_ was shifted by an amount *σ*. We used values of *σ* between −10*T* and 10*T*, at intervals of 0.13*T*. For each such *σ*, the process was simulated 200 times. For each simulation, the TE was estimated on the original process with the shift in the first daughter as well as on a surrogate generated according to our proposed local permutation scheme (see section IV B for a detailed description). The parameter values of *k*_perm_ = 10 and *N_U,_*_surrogate_ = *N_X_* were used. For comparison, we also generated surrogates according to the traditional source time-shift method, where this shift was distributed randomly uniform between 200 and 300. A contiguous region of 50000 target events was extracted and the estimation was performed on this data. The continuous-time estimator used the parameter values of 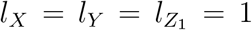, *k* = 10, *N_U_* = *N_X_* and the Manhattan (*ℓ*_1_) norm.

The results in Fig. 6a demonstrate that the null distribution of TE values produced by the the local permutation surrogate generation scheme closely matches the distribution of TE values produced by the continuous-time estimator applied to the original data. Whilst the raw TE estimates retain a slight negative bias (explored further in Discussion), we can generate a bias-corrected TE with the surrogate mean subtracted from the original estimate (giving an “effective transfer entropy” [54]). This bias-corrected TE as displayed in Fig. 6b is consistent with zero because of the close match between our estimated value and surrogates, which is the desired result in this scenario. By contrast, the TE values estimated on the surrogates generated by the traditional time shift method are substantially lower than those estimated on the original process (Fig. 6a); comparison to these would produce very high false positive rates for significant directed relationships. This is most pronounced for high levels of pairwise source-target correlation (with shifts *σ* near zero). The reason behind this difference in the two approaches is easy to intuit. The traditional time-shift method destroys all relationship between the history of the source and the occurrence of events in the target. This means that we are comparing estimates of the TE on processes where there is a strong pairwise correlation between the history of the source and the occurrence of target events, with estimates of the TE on surrogate processes where these are fully independent. Specifically, in discrete time, the joint distribution of the present of the target and the source history, conditioned on the other histories decomposes as 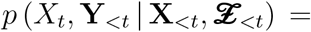 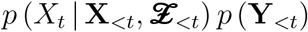.

**FIG. 6:**
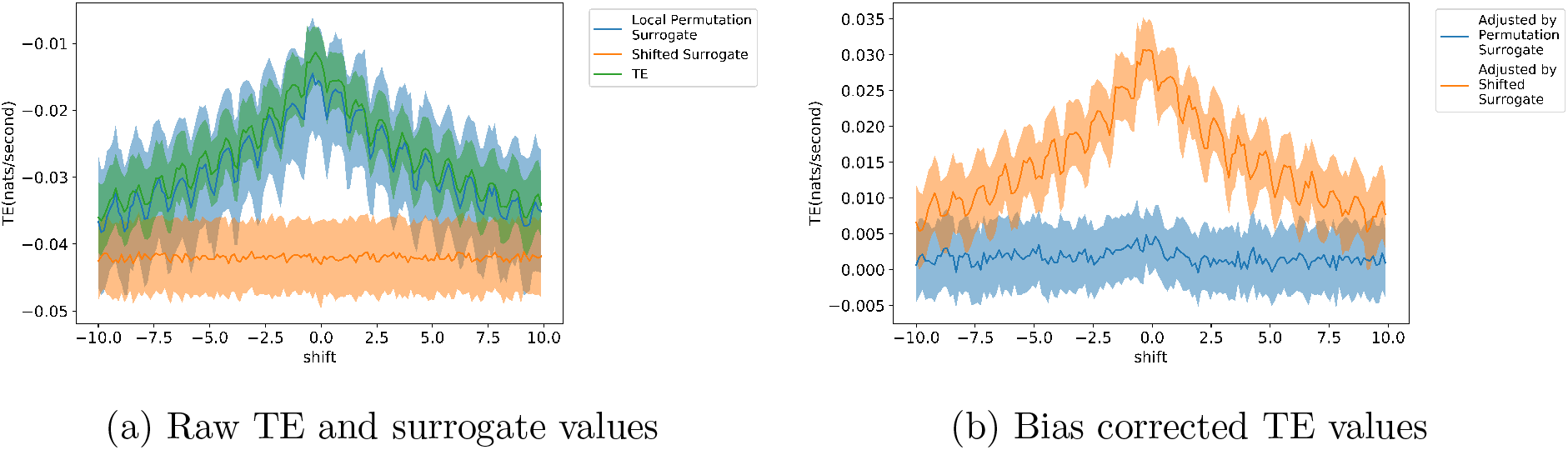
Results of the continuous-time estimator run on a noisy copy process 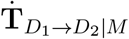, where conditioning on a strong common driver *M* should lead to zero information flow being inferred. The shift *σ* of the source, relative to the target and common driver, controls the strength of the correlation between the source and target (maximal at zero shift). For each shift, the estimator is run on both the original process as well as embeddings generated via two surrogate generation methods: our proposed local permutation method and a traditional source time-shift method. The shaded regions represent the standard deviation. The bias of the estimator changes with the shift, and we expect the estimates to be consistent with appropriately generated surrogates reflecting the same strong common driver effect. This is the case for our local permutation surrogates, as shown in (a). This leads to the correct bias-corrected TE value of 0, as shown in (b).

By contrast, the proposed local permutation scheme produces surrogates where, although the history of the source and the occurrence of events in the target are *conditionally* independent, this scheme maintains the relationship between the history of the source and the mediating variable, which in this case is the history of the mother process. That is, the scheme produces surrogates where (working in discrete time for now) the joint distribution of the present of the target and the source history, conditioned on the other histories decomposes as 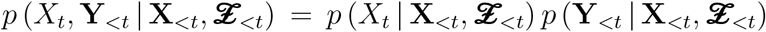. See Methods section IV B for the analogous decomposition within the continuous-time event-based TE framework.

We then confirm that the proposed scheme is able to identify cases where an information flow does exist. To do so, we applied it to measure 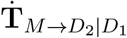 in the above system, where we would expect to see non-zero information flow from the common driver or mother to one daughter process, conditioned on the other. The setup used was identical to above however focussing on a shift of *σ* = 0, and for completeness, two different levels of noise in the daughter processes were used: *σ_D_* = 0.05 and *σ_D_* = 0.075. We recorded the *p* values produced by the combination of the proposed continuous-time estimator and the local permutation surrogate generation scheme when testing for conditional information flow 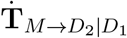, as well as contrasting to the expected zero flow in 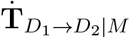. These flows were measured in 10 runs each and the distributions of the resulting *p* values are shown in Fig. 7. We observe that our proposed combination assigns a *p* value of zero in every instance of 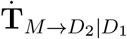 as expected; whilst for 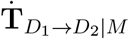 it assigns *p* values in a broad distribution above zero, meaning the estimates are consistent with the null distribution as expected.

**FIG. 7:**
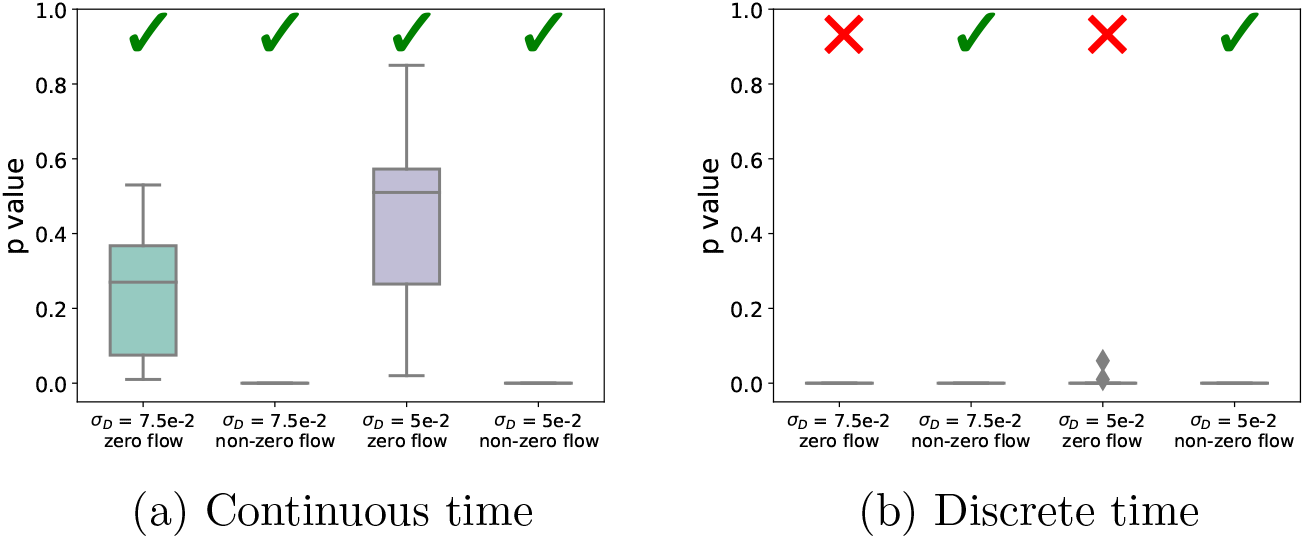
The *p*-values obtained when using continuous and discrete-time estimators to infer non-zero information flow in the noisy copy process. The estimators are applied to both 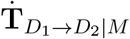 (expected to have zero flow) and 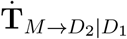 (expected to have non-zero flow and therefore be indicated as statistically significant). Only the results from the continuous-time estimator match these expectations. Ticks represent the particular combination of estimator and surrogate generation scheme making the correct inference in the majority of cases when a cutoff value of *p* = 0.05 is used. Diamonds show outliers.

We also applied the combination of the discrete-time estimator and the traditional time-shift method of surrogate generation to this same task of distinguishing between zero and non-zero conditional information flows. We used time bins of width Δ*t* = 0.05 and history lengths of 7 samples for the target, source and conditional histories. We also applied a procedure to determine optimal lags to the target for both the conditional and source histories, following [17]. That is, before calculating the conditional TE from the source to the target, we determined the optimal lag between the conditional history and the target by calculating the pairwise TE between the conditioning process and the target for all lags between 0 and 10. The lag which produced the maximum such TE was used. We then calculated the conditional TE between the source and the target, using this determined lag for the conditioning process, for all lags to the source process between 0 and 10. The TE was then determined to be the maximum TE estimated over all these different lags applied to the source process. This procedure was applied when estimating the TE on the original process as well as on each separate surrogate. The results of this procedure are also displayed in Fig. 7. Here we see that the combination of the discrete-time estimator and the traditional time-shift method of surrogate generation assigns a *p* value of zero to every single example of both 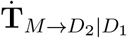 and 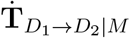. This result – contradicting the expectation that 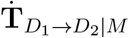 is consistent with zero – suggests that this benchmark approach has an incredibly high false positive rate here.

### D. Connectivity Inference of the Pyloric Circuit of the Crustacean Stomatogastric Ganglion

The pyloric circuit of the crustacean stomatogastric ganglion has been proposed as a benchmark circuit on which to test spike-based connectivity inference techniques [55, 56]. It has been shown that Granger causality (which is equivalent to TE under the assumption of Gaussian variables [57]) is unable to infer the connectivity of this network [55]. In contrast, we demonstrate here the efficacy of the proposed estimators for TE on models of this system.

The crustacean stomatogastric ganglion [58–60] has received substantial research attention as a simple model circuit. Of great importance to network inference is the fact that its full connectivity is known. The pyloric circuit is a partially independent component of the greater circuit and consists of an Anterior Burster (AB) neuron, two Pyloric Driver (PD) neurons, a Lateral Pyloric (LP) neuron and multiple Pyloric (PY) neurons. As the AB neuron is electrically coupled to the PD neurons and the PY neurons are identical, for the purposes of modelling and network inference, the circuit is usually represented by a single AB/PD complex, and single LP and PY neurons [55, 56, 61, 62].

The AB/PD complex undergoes self-sustained rhythmic bursting. It inhibits the LP and PY neurons through slow cholinergic and fast glutamatergic synapses. These neurons then burst on rebound from this inhibition. The PY and LP neurons also inhibit one another through fast glutamatergic synapses and the LP neuron similarly inhibits the AB/PD complex.

Fig. 8 shows sample membrane potential traces from simulations of this circuit as well as a connectivity diagram. Despite its small size, inference of its connectivity is challenging [55, 56] due to the fact that it is highly periodic. Although there is no structural connection from the PY to the ABPD neuron, there is a strong, time-directed, correlation between their activity — the PY neuron always bursts shortly before the ABPD. Recognising that this is a spurious correlation requires fully resolving the influence of the AB/PD’s history on itself as well as that of the LP on the AB/PD. To further complicate matters, the true connection between the LP and ABPD neurons is very challenging to detect. The AB/PD complex will continue bursting regardless of any input from the LP. Correctly inferring this connection requires detecting the subtle changes in the timing of AB/PD bursts that result from the activity of the LP.

**FIG. 8:**
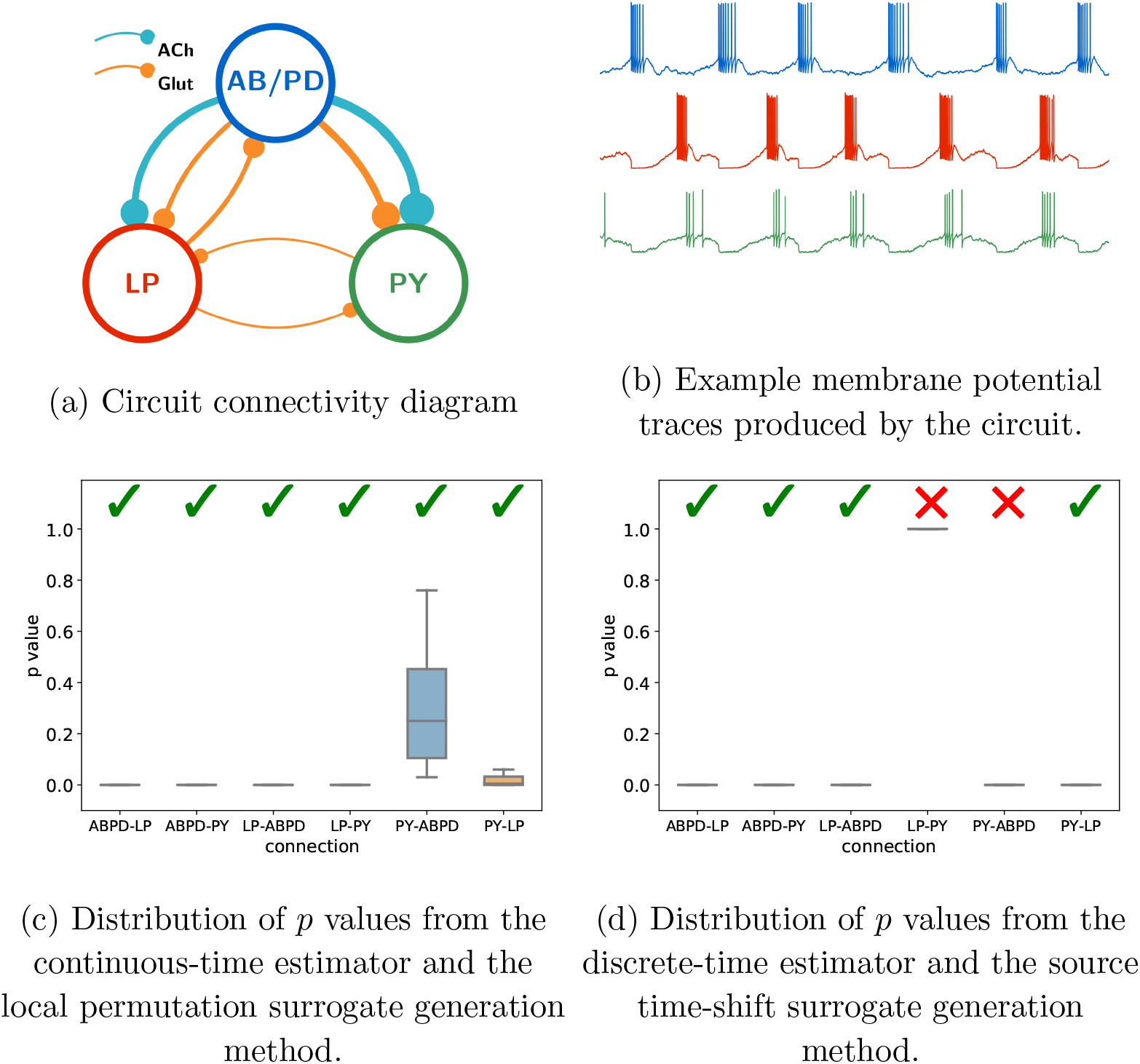
Results of both estimator and surrogate generation combinations being applied to data from simulations of a biophysical model of a neural circuit inspired by the pyloric circuit of the crustacean stomatogastric ganglion. The circuit, shown in (a), is fully connected apart from the missing connection between the PY neuron and the AB/PD complex, and generates membrane potential traces which are bursty and highly-periodic with cross-correlated activity. The distribution of *p* values from the combination of the continuous-time estimator and local permutation surrogate generation scheme shown in (c) demonstrates that this combination is capable of correctly inferring this circuit in most runs. By contrast, the distribution of *p* values produced by the combination of the discrete-time estimator and the traditional source time-shift surrogate generation method shown in (d) mis-specified the connectivity in every run. Ticks represent the particular combination of estimator and surrogate generation scheme making the correct inference in the majority of cases when a cutoff value of *p* = 0.05 is used.

Previous work on the inference of the pyloric circuit has used both *in vitro* and *in silico* data [55, 56]. We ran simulations of biophysical models inspired by this network, similar to those used in [55] (see Appendix A). Attempts were then made to infer this network by detecting non-zero conditional information flow from the spiking event times produced by the simulations. This was done by, for every source-target pair, estimating the TE from the source to the target, conditioned on the activity of the third remaining neuron. Both the combination of the proposed continuous-time estimator and local permutation surrogate generation scheme and the combination of the discrete-time estimator and source time-shift surrogate generation scheme were applied to this task.

Both combinations were applied to ten independent simulations of the network and the number of target events *N_X_* = 1 × 10^4^ was used. For the continuous-time estimator the parameter values of 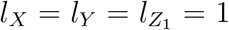, *k* = 10, *N_U_* = *N_X_*, *N_U,_*_surrogate_ = *N_X_* and *k*_perm_ = 5 were used along with the Manhattan (*ℓ*_1_) norm (see Methods Sec. IV A 4 and IV A 5). The discrete-time estimator made use of a bin size of Δ*t* = 0.05 and history embedding lengths of seven bins for each of the source, target and conditioning processes. Searches were performed to determine the optimum embedding lag for both the source and conditioning histories (as per Sec. II C) with a maximum search value of 20 bins being used. We designed the search procedure to include times up to the inter-burst interval (around 1 time unit), which placed an effective lower bound on the width of the time bins (as bin sizes below Δ*t* = 0.05 resulted in impractically large search spaces). For both estimators, *p* values were inferred from 100 independently generated surrogates (see Methods Sec. IV B). The source time-shift surrogate generation scheme used time shifts distributed uniformly randomly between 200 and 400 time units.

Figures 8c and 8d show the distributions of *p* values resulting from the application of both estimator and surrogate generation scheme combinations. The continuous-time estimator and local permutation surrogate generation scheme were able to correctly infer the connectivity of the network in the majority of cases (indicated by *p*-values approaching 0 for the true positives, and spread throughout [0,1] for the true negatives). On the other hand, the discrete-time estimator and source time-shift surrogate generation scheme produced the same two erroneous predictions on every run: a connection between the PY neuron and the AB/PD complex along with a missing connection between the LP and PY neurons.

## III. DISCUSSION

Despite transfer entropy being a popular tool within neuroscience and other domains of enquiry [9–11, 13–19], it has received more limited application to event-based data such as spike trains. This is at least partially due to current estimation techniques requiring the process to be recast as a discrete-time phenomenon. The resulting discrete-time estimation task has been beset by difficulties including a lack of consistency, high bias, slow convergence and an inability to capture effects which occur over fine and large time scales simultaneously.

This paper has built on recent work presenting a continuous time formalism for TE [5] in order to derive an estimation framework for TE on event-based data in continuous time. This framework has the unique advantage of only estimating quantities at events in the target process, providing a significant computational advantage. Instead of comparing spike rates conditioned on specific histories at each target spiking event though, we use a Bayesian inversion to instead make the empirically easier comparison of probabilities of histories at target events versus anywhere else along the target process. This comparison, using KL divergences, is made using *k*-NN techniques, which brings desirable properties such as efficiency for the estimator. This estimator is provably consistent. Moreover, as it operates on inter-event intervals, it is capable of capturing relationships which occur with fine time precision along with those that occur over longer time distances.

The estimator was first evaluated on two simple examples for which the ground truth is known: pairs of independent Poisson processes (Sec. II A) as well as pairs of processes unidirectionally coupled through a simple functional relationship (Sec. II B). The discrete-time estimator was also applied to these processes. It was found that the continuous-time estimator had substantially lower bias than the discrete-time estimator, converged orders of magnitude faster (in terms of the number of sample spikes required), and was relatively insensitive to parameter selections. Moreover, these examples provided numerical confirmation of the consistency of the continuous-time estimator, and further demonstration that the discrete-time estimator is not consistent. The latter simple example highlighted the magnitude of the shortcomings of the discrete-time estimator. In the authors’ experience, spike-train datasets which contain 1 million spiking events for a single neuron are vanishingly rare. However, even in the unlikely circumstance that the discrete-time estimator is presented with a dataset of this size as in Sec. II B, it could not accurately estimate the TE for a simple one-way relationship between only two neurons. Moreover, this example neatly demonstrates a notable problem with the use of the discrete-time estimator, which is that it provides wildly different estimates for different values of Δ*t*. In real-world applications, where the ground truth is unknown, there is no principled method for choosing which resulting TE value from the various bin sizes to use.

One of the principal use-cases of TE is the inference of non-zero information flow. As the TE is estimated from finite data, we require a manner of determining the statistical significance of the estimated values. Traditional methods of surrogate generation for TE either shift the source in time, or shuffle the source embeddings. However, whilst this retains the relationship of the target to its past and other conditionals, it completely destroys the relationship between the source and any conditioning processes, which can lead to very high false positive rates as detailed in Results Sec. II C and Sec. IV B. We developed a local permutation scheme for use in conjunction with this estimator which is able to maintain the relationship of the source history embeddings with the history embeddings of the target and conditioning processes. The combination of the proposed estimator and this surrogate generation scheme were applied to an example where the history of the source and the occurrence of events in the target are highly correlated, but conditionally independent given their common driver (Sec. II C). The established time-shift method for surrogate generation produced a null distribution of TE values substantially below that estimated on the original data, incorrectly implying non-zero information flow. Conversely, the proposed local permutation method produced a null distribution which closely tracked the estimates on the original data. The proposed combination was also shown to be able to correctly distinguish between cases of zero and non-zero information flow. When applied to the same example, the combination of the discrete-time estimator and the traditional method of time-shifted surrogates inferred the existence of information flow in all cases, even when no such flow was present.

Finally, our proposed approach was applied to the inference of the connectivity of the pyloric network of the crustacean stomatogastric ganglion. The inference of this network is challenging due to highly periodic dynamics. Granger causality, using a more established estimator, has historically been demonstrated to be incapable of inferring its connectivity [55]; furthermore, we showed that the discrete-time binary-valued TE estimator (with time-shifted surrogates) also could not successfully infer the connectivity here. Despite these challenges, our combination of continuous-time estimator and surrogate generation scheme was able to correctly infer the connectivity of this network. This provides an important validation of the efficacy of our presented approaches on a challenging example of representative biological spiking data.

This work represents a substantial step forward in the estimation of information flows from event-based data. To the best of the authors’ knowledge it is the first consistent estimator of TE for event-based data. That is, it is the first estimator which is known to converge to the true value of the TE in the limit of infinite data, let alone to provide efficient estimates with finite data. As demonstrated in Results Sec. II A and Sec. II B it has substantially favourable bias and convergence properties as compared to the discrete-time estimator. The fact that this estimator uses raw inter-event intervals as its history representation allows it to efficiently capture relevant information from the past of the source, target and conditional processes. This allows it to simultaneously measure relationships that occur both with very fine time scales as well as those that occur over long intervals. This was highlighted in Results Sec. II D, where it was shown that our proposed approach is able to correctly infer the connectivity of the pyloric circuit of the crustacean stomatogastric ganglion. The inference of this circuit requires capturing subtle changes in spike timing. However, its bursty nature means that there are long intervals of no spiking activity. This is contrasted with the poor performance of the discrete-time estimator on this same task, as above. The use of the discrete-time estimator requires a hard trade-off in the choice of bin size: small bins will be able to capture relationships that occur over finer timescales but will result in an estimator that is blind to history effects existing over large intervals. Conversely, whilst larger bins might be capable of capturing these relationships occurring over larger intervals, the estimator will be blind to effects occurring with fine temporal precision. Moreover, to the best of our knowledge, this work showcases the first use of a surrogate generation scheme for statistical significance estimates which correctly handles strong source-conditional relationships for event-based data. This has crucial practical benefit in that it greatly reduces the occurrence of false positives in cases where the history of a source is strongly correlated with the present of the target, but conditionally independent.

Inspection of some plots, notably Fig. 6 of Sec. II C shows that, in some cases, the estimator can exhibit small though non-insignificant bias. Indeed, similar biases can readily be demonstrated with the standard KSG estimator for transfer entropy on continuous variables in discrete time, in similar circumstances where a strong source-target relationship is fully explained by a conditional. The reason for the small remaining bias is that while underlying assumption of the nearest neighbour estimators is of a fixed probability density within the range of the *k* nearest neighbours, strong conditional relationships tend to result in correlations remaining between the variables within this range. For the common use-case of inferring non-zero information flows this small remaining bias will not be an issue as the proposed method for surrogate generation is capable of producing null distributions with very similar bias properties. Furthermore, such bias can be removed from an estimate by subtracting the mean of the surrogate distribution (as shown via the effective transfer entropy [54] in Results). However, it is foreseeable that certain scenarios might benefit from an estimator with lower bias, without having to resort to generating surrogates. In such cases it will likely prove beneficial to explore the combination of various existing bias reduction techniques for *k*-NN estimators with the approach proposed here. These include performing a whitening transformation on the data [63], transforming each marginal distribution to uniform or exploring alternative approaches to sharing radii across entropy terms (see Sec. IV A 5). The authors believe that the most probable cause of the observed bias in the case of strong pairwise correlations is that it causes the assumption of local uniformity (see Methods) to be violated. Gao et. al [64] have proposed a method for reducing the bias of *k*-NN information theoretic estimators which specifically addresses cases where local uniformity does not apply. The application of this technique to our estimator holds promise for addressing this bias.

## IV. METHODS

There are a variety of approaches available for estimating information theoretic quantities from continuous-valued data [65]; here we focus on methods for generating estimates 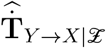 of a true underlying (conditional) transfer entropy 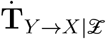.

The nature of estimation means that our estimates 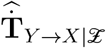 may have a *bias* with respect to the true value 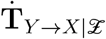, and a *variance*, as a function of some metric *n* of the size of the data being provided to the estimator (we use the number of spikes, or events, in the target process). The bias is a measure of the degree to which the estimator systematically deviates from the true value of the quantity being estimated, for finite data size. It is expressed as bias 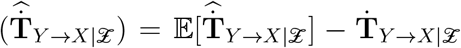. The variance of an estimator is a measure of the degree to which it provides different estimates for distinct, finite, samples from the same process. It is expressed as variance 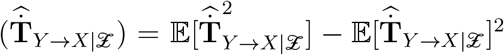. Another important property is *consistency*, which refers to whether, in the limit of infinite data, the estimator converges to the true value. That is, an estimator is consistent if and only if 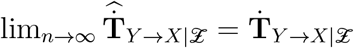.

The first half of this methods section is concerned with the derivation of a consistent estimator of TE which operates in continuous time. In order to be able to test for non-zero information flow given finite data, we require a surrogate generation scheme to use in conjunction with the estimator. Such a surrogate generation scheme should produce surrogate history samples that conform to the null hypothesis of zero information flow. The second half of this section will focus on a scheme for generating these surrogates.

The presented estimator and surrogate generation scheme have been implemented in a software package which will be made available on publication (see the Implementation subsection).

### A. Continuous-time estimator for transfer entropy between spike trains

In the following subsections, we describe the algorithm for our estimator 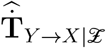 for the transfer entropy between spike trains. We first outline our choice of a *k*NN type estimator, due to the desirable consistency and bias properties of this class of estimator. In order to use such an estimator type, we then describe a Bayesian inversion we apply to the definition of transfer entropy for spiking processes, which allows us to operate on probability densities of histories of the processes, rather than directly on spike rates. This results in a sum of differential entropies to which *k*NN estimator techniques can be applied. The evaluation of these entropy terms using *k*NN estimators requires a method for sampling history embeddings, which is presented before attention is turned to a technique for combining the separate *k*NN estimators in a manner that will reduce the bias of the final estimate.

#### 1. Consideration of estimator type

Although there has been much recent progress on parametric information-theoretic estimators [66], such estimators will always inject modelling assumptions into the estimation process. Even in the case that large, general, parametric models are used — as in [67] — there are no known methods of determining whether such a model is capturing all dependencies present within the data.

In comparison, nonparametric estimators make less explicit model assumptions regarding the probability distributions. Early approaches included the use of kernels for the estimation of the probability densities [68], however this has the disadvantage of operating at a fixed kernel ‘resolution’. An improvement was achieved by the successful, widely applied, class of nonparametric estimators making use of *k*-nearest-neighbour statistics [44, 69–71], which dynamically adjust their resolution given the local density of points. Crucially, there are consistency proofs [70, 72] for *k*NN estimators, meaning that these methods are known to converge to the true values in the limit of infinite data size. These estimators operate by decomposing the information quantity of interest into a sum of differential entropy terms *H*_∗_. Each entropy term is subsequently estimated by estimating the probability densities *p*(*x_i_*) at all the points in the sample by finding the distances to the *k*th nearest neighbours of the points *x_i_*. The average of the logarithms of these densities is found and is adjusted by bias correction terms. In some instances, most notably the KSG estimator for mutual information [44], many of the terms in each entropy estimate cancel and so each entropy is only implicitly estimated.

Such bias and consistency properties are highly desirable – given the efficacy of *k*NN estimators, it would be advantageous to be able to make use of such techniques in order to estimate the transfer entropy of point processes in continuous time. However the continuous time formulations in (3) and (4) contain no entropy terms, being written in terms of *rates* as opposed to probability densities. Moreover, the estimators for each differential entropy term *H*_∗_ in a standard *k*NN approach operate on sets of points in ℝ^*d*^, and it is unclear how these may be applied in our case.

The following sub-section is concerned with deriving an expression for continuous-time transfer entropy on spike trains as a sum of *H*_∗_ terms, in order to define a *k*NN type estimator.

#### 2. Formulating continuous-time TE as a sum of differential entropies

Consider two point processes *X* and *Y* represented by sets of real numbers, where each element represents the time of an event. That is, 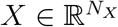 and 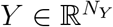. Further, consider the set of extra conditioning point processes 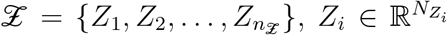. We can define a *counting process N_X_*(*t*) on *X*. *N_X_*(*t*) is a natural number representing the ‘state’ of the process. This state is incremented by one at the occurrence of an event. The instantaneous firing rate of the target is then *λ_X_*(*t*) = lim_Δ*t*→0_ *p* (*N_X_*(*t* + Δ*t*) − *N_X_*(*t*) = 1) /Δ*t*. Using this expression, (4) can then be rewritten as

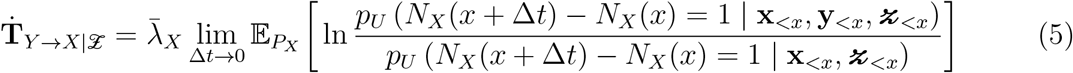

Here, 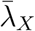 is the average, unconditional, firing rate of the target process 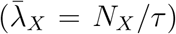, whilst **x**_<*x*_ ∈ **X**_<*X*_, **y**_<*x*_ ∈ **Y**_<*X*_ and 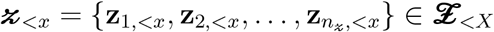 are the histories of the target, source and conditioning processes, respectively, at time *x*. The probability density *p_U_* is taken to represent the probability density at any arbitrary point in the target process, unconditional of events in any of the processes. By contrast, *p_X_* is taken to represent the probability density of observing a quantity at target events. The expectation 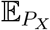 is taken over this distribution. That is 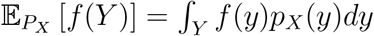

By applying Bayes’ rule we can make a Bayesian inversion to arrive at:

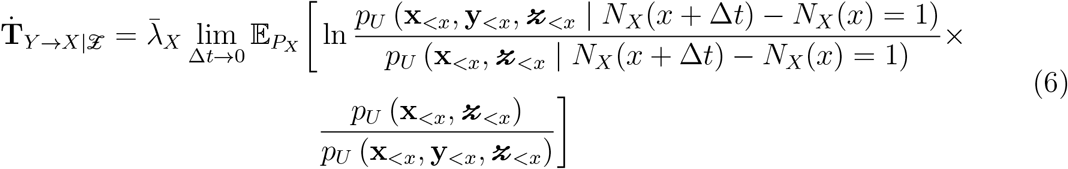

Equation (6) can be written as

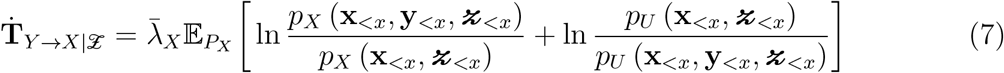

Equation (7) can be written as a sum of differential entropy and cross entropy terms.

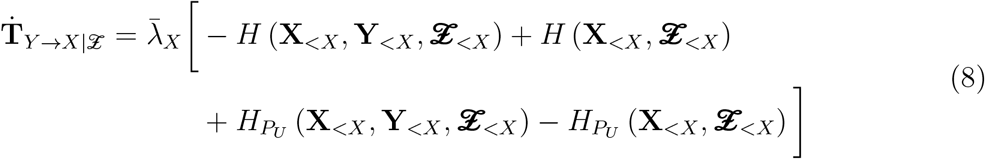

Here, *H* refers to an entropy term and 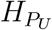 refers to a cross entropy term. More specifically,

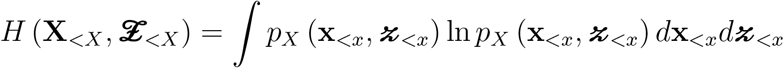

and

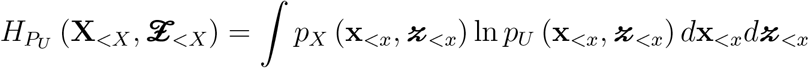

It is worth noting in passing that (7) can also be written as a difference of Kullback-Leibler divergences:

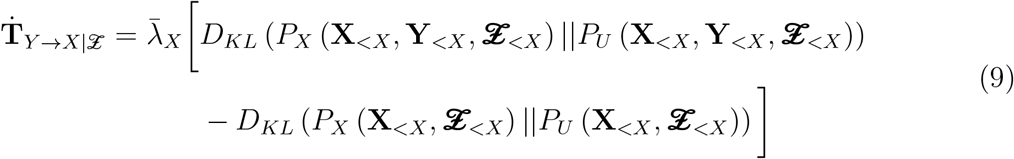

The expressions in equations (8) and ((9) represent a general framework for estimating the TE between point processes in continuous time. Any estimator of differential entropy 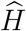 which can be adapted to the estimation of cross entropies can be plugged into (8) in order to estimate the TE. Similarly, any estimator of the KL divergence can be plugged into (9).

#### 3. Constructing kNN estimators for differential entropies and cross entropies

Following similar steps to the derivations in [44, 63, 72], assume that we have an (unknown) probability distribution *μ*(**x**) for **x**∈ ℝ*^d^*. Note that here **X** is a general random variable (not necessarily a point process). We also have a set *X* of *N_X_* points drawn from *μ*.

In order to estimate the differential entropy *H* we need to construct estimates of the form

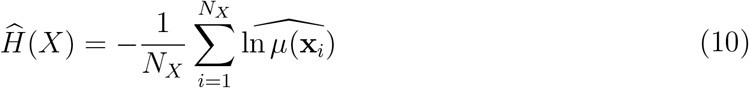

where 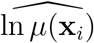 is an estimate of the logarithm of the true density. Denote by *ϵ* (*k,* **x**_*i*_, _*X*_) the distance to the *k*th nearest neighbour of **x**_*i*_ in the set *X* under some norm *L*. Further, let 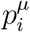 be the probability mass of the *ϵ*-ball surrounding **x**_*i*_. If we make the assumption that *μ*(**x**_*i*_) is constant within the *ϵ*-ball, we have 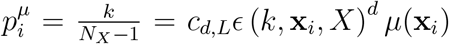 where *c_d,L_* is the volume of the *d*-dimensional unit ball under the norm *L*. Using this relationship, we can construct a simple estimator of the differential entropy:

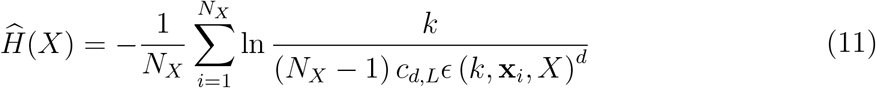

We then add the bias-correction term ln *k* − *ψ*(*k*). *ψ*(*x*) = Γ^−1^(*x*)*d*Γ(*x*)*/dx* is the digamma function and Γ(*x*) the gamma function. This yields 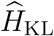, the Kozachenko-Leonenko [69] estimator of differential entropy:

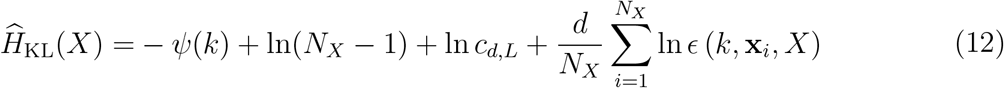

This estimator has been shown to be consistent [69, 73].

Assume that we now have two (unknown) probability distributions *μ*(**x**) and *β*(**x**). We have a set *X* of *N_X_* points drawn from *μ* and a set *Y* of *N_Y_* points drawn from *β*. Using similar arguments to above, we denote by *ϵ* (*k,* **x**_*i*_, *Y*) the distance from the *i*th element of *X* to its *k*th nearest neighbour in *Y*. We then make the assumption that *β*(**x**_*i*_) is constant within the *ϵ*-ball, and we have 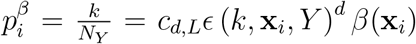. We can then construct a naive estimator of the cross entropy

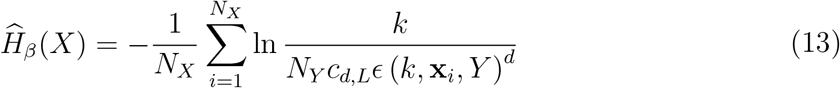

Again, we add the bias-correction term ln *k* − *ψ*(*k*) to arrive at an estimator of the cross entropy.

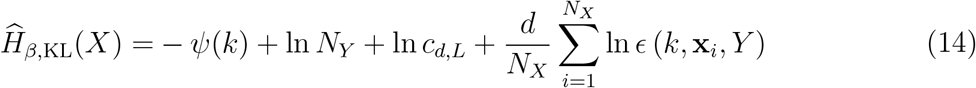

This estimator has been shown to be consistent [73].

Attention should be brought to the fundamental difference between estimating entropies and cross entropies using *k*-NN estimators. An entropy estimator takes a set *X* and, for each *x_i_* ∈ *X*, performs a nearest neighbour search *in the same set X*. An estimator of cross entropy takes two sets, *X* and *Y* and, for each *x_i_* ∈ *X*, performs a nearest neighbour search *in the other set Y*.

We will be interested in applying these estimators to the entropy and cross entropy terms in (8). For instance, we could use 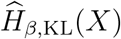 to estimate 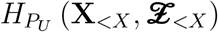, where we have that 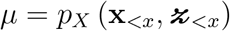 and 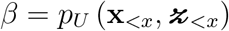. This will be covered in more detail in section IV A 5, after we first consider how to represent the history embeddings 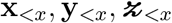 as well as sample them from their distributions.

#### 4. Selection and Representation of Sample Histories for Entropy Estimation

Inspection of (7) and (8) informs us that we will need to be able to estimate four distinct differential entropy terms and, implicitly, the associated probability densities:

1. The probability density of the target, source and conditioning histories at target events 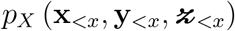.
2. The probability density of the target, and conditioning histories at target events 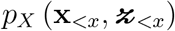.
3. The probability density of the target, source and conditioning histories independent of target activity 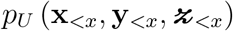.
4. The probability density of the target and conditioning histories independent of target activity 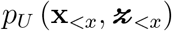.

##### Algorithm 1: the CT TE estimator

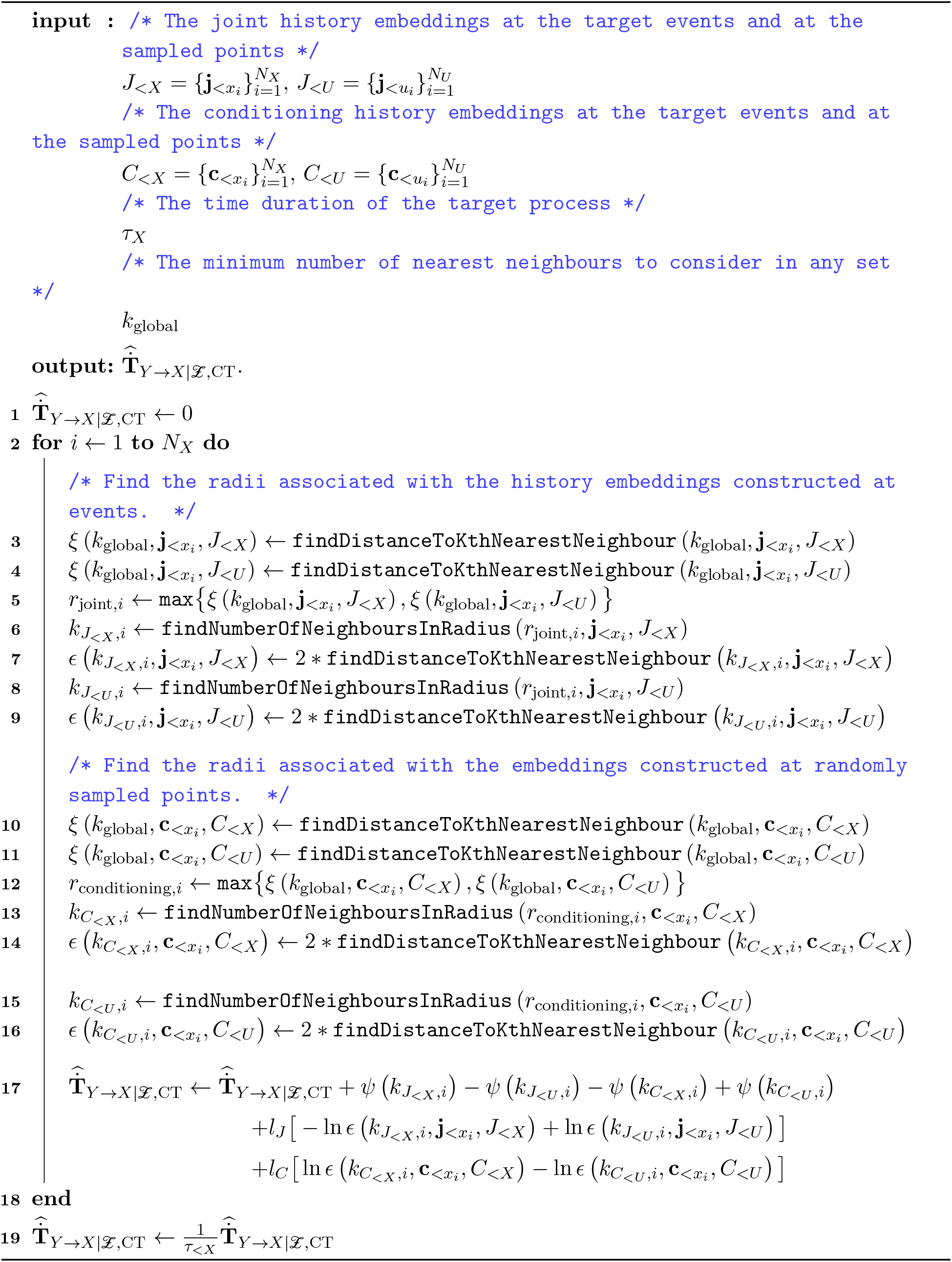

Estimation of these probability densities will require an associated set of samples for a *k*-NN estimator to operate on. These samples for 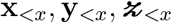 will logically be representated as history embeddings from the raw event times of the target 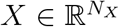, source 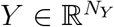 and conditioning 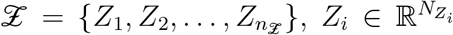 processes. It is assumed that these sets are indexed in ascending order (from the first event to the last). The length of the history embeddings (in terms of how many previous spikes are referred to) must be restricted in order to avoid the difficulties associated with the estimation of probability densities in high dimensions. The lengths of the history embeddings along each process are specified by the parameters 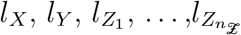.

We label the sets of samples as 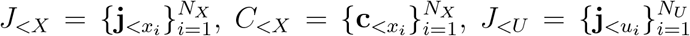, and 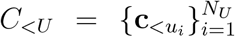, for each probability density 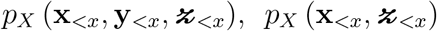, 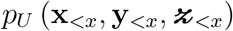 and 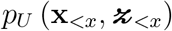 respectively (*J* for ‘joint’ and *C* for ‘conditioning’, i.e. without the source).

For the two sets of joint embeddings *J*_<∗_ (where ∗ ∈ {*X, U*}) each 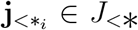 is made up of target, source and conditioning components. That is, 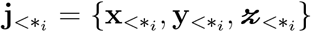 where 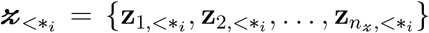. Similarly, for the two sets of conditioning embeddings *C*_<∗_ (where ∗ ∈ {*X, U*}) each 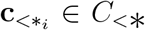 is made up of target, and conditioning components. That is, 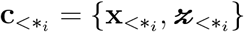.

Each set of embeddings *J*_<∗_ is constructed from a set of observation points 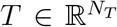. Each individual embedding 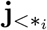 is constructed at one such observation *t_i_*. We denote by pred (*t_i_, P*), the index of the most recent event in the process *P* to occur before the observation point *t_i_*.

The values of 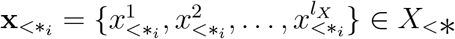 are set as follows:

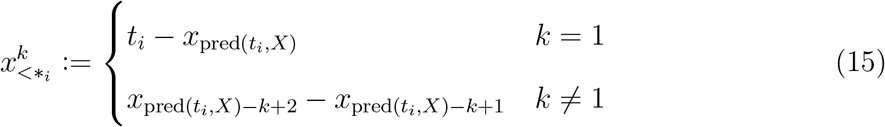

Here, the *t_i_* ∈ *T* are the raw observation points and the *x_j_* ∈ *X* are the raw event times in the process *X*. The first element of 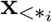 is then the interval between the observation time and the most recent target event time 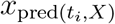. The second element of 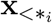 is the inter-event interval between this most recent event time and the next most recent event time and so forth. The values of 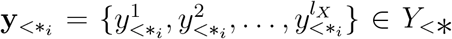 and 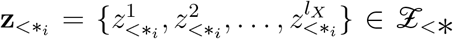 are set in the same manner.

The set of samples 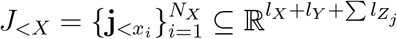 for 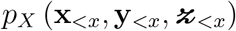 is constructed using this scheme, with the set of observation points *T* being simply set as the *N_X_* event times *x_j_* of the target process *X*. As such, 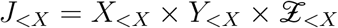.

In contrast, while the set of samples 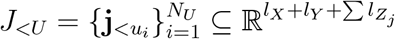 for 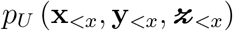 is also constructed using this scheme, the set of observation points *T* is set as 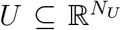. *U* is composed of sample time points placed independently of the occurrence of events in the target process. These *N_U_* sample points between the first and last events of the target process *X* can either be placed randomly or at fixed intervals. In the experiments presented in this paper they were placed randomly. Importantly, note that *N_U_* is not necessarily equal to *N_X_*, with their ratio *N_U_ /N_X_* a parameter for the estimator which is investigated in our Results. We also have that 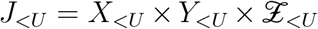. Figure 9 shows diagramatic examples of an embedded sample from *J*_<*X*_ as well as one from *J_<U_*. Notice the distinction that for *J*_<*X*_, the 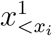 in the embeddings 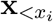 are specifically an interspike interval from the current spike at *t_i_* = *x_i_* back to the previous spike, which is not the case for *J_<U_*.

**FIG. 9:**
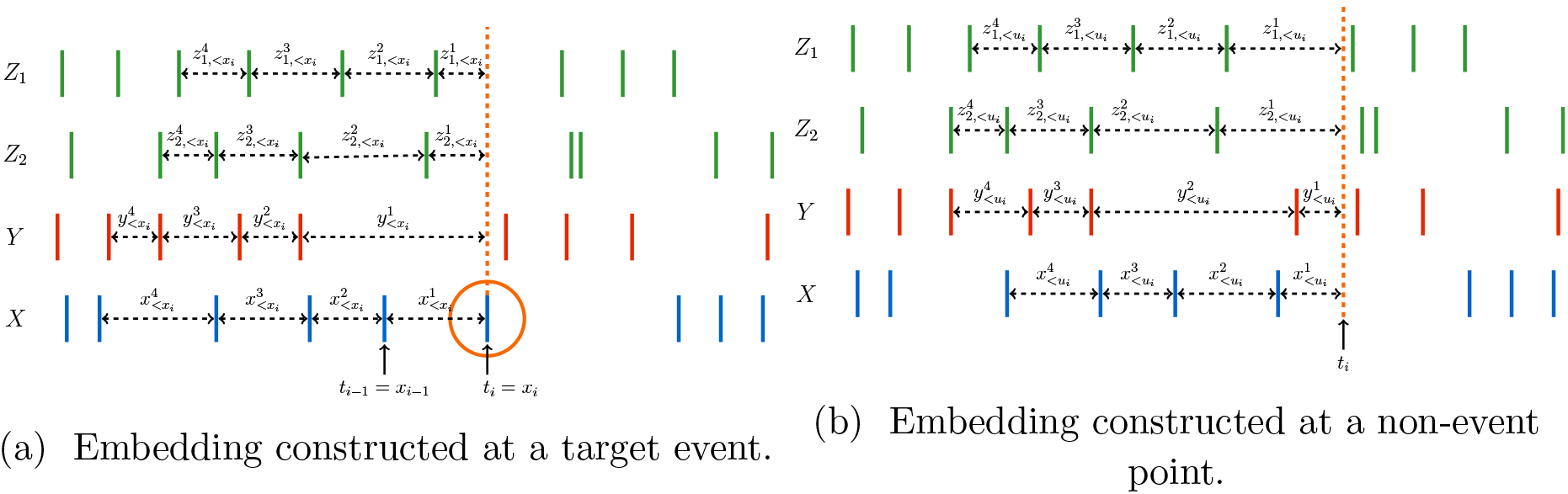
Examples of history embeddings. (a) shows an example of a joint embedding constructed at a target event 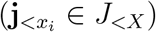. (b) shows an example of a joint embedding constructed at a sample event 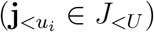.

The set of samples 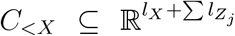 for 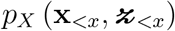 and 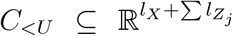 for 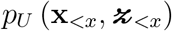 are constructed in a similar manner to their associated sets *J_<X_* and *J*_<*U*_, how-ever, the source embeddings 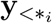 are discarded. We will also have that 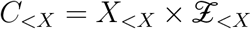 and 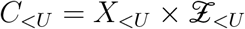.

Note that, as 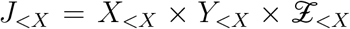 and 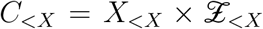, these two sets are closely related. Specifically, the *i*-th element of *C*_<*X*_ will be identical to the *i*-th element of *J*_<*X*_, apart from missing the source component 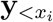.

#### 5. Combining 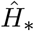 estimators for 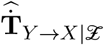

With sets of samples and their embedded representation determined as per the previous subsection, we are now ready to estimate each of the four 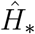 terms in (8). Here we consider how to combine the entropy and cross entropy estimators of these terms ((12) and (14)) into a single estimator.

We could simply estimate each *H*_∗_ term in (8) using 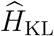 as specified in (12) and 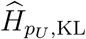 as specified in (14), with the same number *k* of nearest neighbours in each of the four estimators and at each sample in the set for each estimator. Following the convention introduced in [72] we shall refer to this as a 4KL estimator of transfer entropy (the ‘4’ refers to the use 4 *k*-NN searches and the ‘KL’ to Kozachenko-Leonenko):

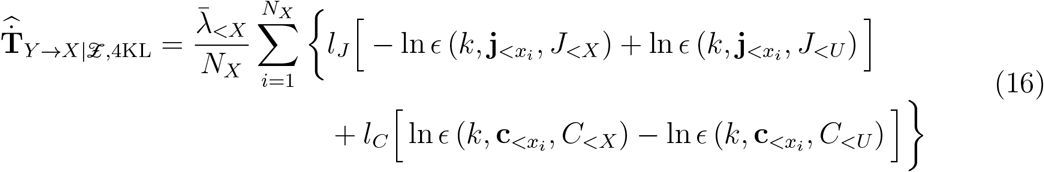

Here, 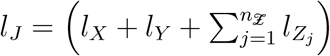 is the dimension of the joint samples and 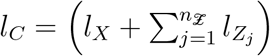 is the dimension of the conditional-only samples. Note that the ln(*N_X_* − 1) − *ψ*(*k*) terms cancel between the *J*_<*X*_ and *C*_<*X*_ terms (also for ln(*N_U_*) − *ψ*(*k*) between the *J_<U_* and *C_<U_* terms), whilst the ln *c_d,L_* terms cancel between *J*_<*X*_ and *J_<U_* as well as between *C*_<*X*_ and *C_<U_*. It is crucial also to notice that all terms are averaged over *N_X_* samples taken at target events (the cross-entropies which evaluate probability densities using *J_<U_* and *C_<U_* still evaluate those densities on the samples 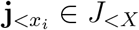 and 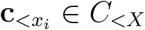, following the definition in (14)), regardless of whether *N_U_* = *N_X_*.

It is, however, not only possible to use a different *k* at every sample, but desirable when the *k* are chosen judiciously (as detailed below). We shall refer to this as the *generalised k*-NN estimator:

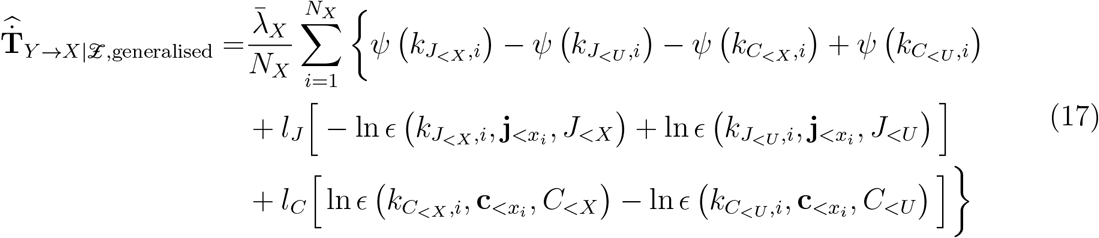

Here *k_A,i_* is the number of neighbours used for the *i*th sample in set *A* for the corresponding entropy estimator for that set of samples. By theorems 3 and 4 of [63] this estimator (and, by implication, the 4KL estimator) is asymptotically consistent. Application of the generalised estimator requires a scheme for choosing the *k_A,i_* at each sample. Work on constructing *H*_∗_ *k*-NN estimators for mutual information [44] and KL divergence [63] has found advantages in having certain *H*_∗_ terms share the same or similar radii, e.g. resulting in lower overall bias due to components of biases of individual *H*_∗_ terms cancelling. Given that we have four *H*_∗_ terms, there are a number of approaches we could take to sharing radii.

Our algorithm, which we refer to as the CT estimator of 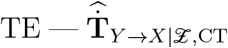 is specified in detail in algorithm 1. Our algorithm applies the approach of Wang et. al. (referred to as the ‘bias improved’ estimator in [63]) to each of the Kullback-Leibler divergence terms separately. In broad strokes, whereas, (16) uses the same *k* for each nearest-neighbour search, this estimator uses the same *radius* for each of the two nearest-neighbour searches relating to a given KL divergence term. In practice, this requires first performing searches with a fixed *k* in order to determine the radius to use. As such, we start with a fixed parameter *k*_global_, which will be the minimum number of nearest neighbours in any search space. For each joint sample at a target event, that is, each 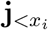 in *J*_<*X*_, we perform a *k*_global_-NN search in this same set *J*_<*X*_ and record the distance to the *k*_global_-th nearest neighbour (line 3 of algorithm 1). We perform a similar *k*_global_-NN search for 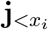 in the set of joint samples independent of target activity *J_<U_*, again recording the distance to the *k*_global_-th nearest neighbour (line 4). We define a search radius as the maximum of these two distances (line 5). We then find the number of points in *J*_<*X*_ that fall within this radius of 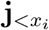 and set 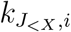 as this number (line 6). We also find twice the distance to the 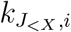-th nearest neighbour, which is the term 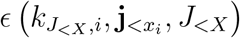 in (17) (line 7). Similarly, we find the number of points in *J_<U_* that fall within the search radius of 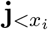 and set 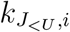 as this number (line 8). We find twice the distance to the 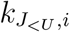-th nearest neighbour, which is the term 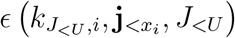 (line 9).

In the majority of cases, only one of these two *ϵ* terms will be exactly twice the search radius, and its associated *k_A,i_* will equal *k*_global_. In such cases, the other *ϵ* will be less than twice the search radius and its associated *k_A,i_* will be greater than or equal to *k*_global_.

The same set of steps is followed for each conditioning history embedding that was constructed at an event in the target process, that is, each 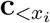 in *C*_<*X*_, over the sets *C*_<*X*_ and *C*_<*U*_ (lines 10 through 16 of algorithm 1).

The values that we have found for 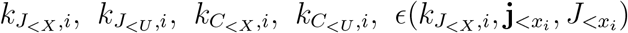, 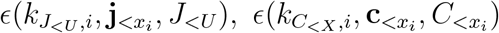 and 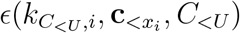 can be plugged into (17) (lines 17 and 19 of algorithm 1).

#### 6. Handling Dynamic Correlations

The derivation of the *k*-NN estimators for entropy and cross entropy given in section IV A 3 assumes that the points are independent [44] However, nearby interspike intervals might be autocorrelated (e.g. during bursts), and indeed our method for constructing history embeddings (see section IV A 4) will incorporate the same interspike intervals at different positions in consecutive samples. This contradicts the assumption of independence. In order to satisfy the assumption of independence when counting neighbours, conventional neighbour counting estimators can be made to ignore matches within a dynamic or serial correlation exclusion window (a.k.a. Theiler windows [74, 75]).

For our estimator, we maintain a record of the start and end times of each history embedding, providing us with an exclusion window. The start time is recorded as the time of the first event that formed part of an interval which was included in the sample. This event could come from the embedding of any of the processes from which the sample was constructed. The end of the window is the observation point from which the sample is constructed. When performing nearest neighbour and radius searches (lines 3, 4, 6, 7, 8, 9, 10, 11, 13, 14, 15 and 16 of algorithm 1 and line 6 of algorithm 2), any sample whose exclusion window overlaps with the exclusion window of the sample around which the search is taking place is ignored. Subtleties concerning dynamic correlation exclusion for surrogate calculations are considered in the next section.

### B. Local Permutation Method for Surrogate Generation

A common use of this estimator would be to ascertain whether there is a non-zero conditional information flow between two components of a system. When using TE for directed functional network inference, this is the criteria we use to determine the presence or absence of a connection. Given that we are estimating the TE from finite samples, we require a statistical test in order to determine the significance of the measured TE value. Unfortunately, analytic results do not exist for the sampling distribution of *k*-NN estimators of information theoretic quantities [8]. This necessitates a scheme for generating surrogate samples from which the null distribution can be empirically constructed.

It is instructive to first consider the more general case of testing for non-zero mutual information. As the mutual information between *X* and *Y* is zero if and only *X* and *Y* are independent, testing for non-zero mutual information is a test for statistical dependence. As such, we are testing against the null hypothesis that *X* and *Y* are independent (*X* ⫫ *Y*) or, equivalently, that the joint probability distribution of *X* and *Y* factorises as *p*(*X, Y*) = *p*(*X*)*p*(*Y*). It is straightforward to construct surrogate pairs 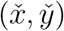 that conform to this null hypothesis. We start with the original pairs (*x, y*) and resample the *y* values across pairs, commonly by shuffling (in conjunction with handling dynamic correlations, as per the later subsection). This shuffling process will maintain the marginal distributions *p*(*X*) and *p*(*Y*), and the same number of samples, but will destroy any relationship between *X* and *Y*, yielding the required factorisation for the null hypothesis. One shuffling process produces one set of surrogate samples; estimates of mutual information on populations of such surrogate sample sets yields a null distribution for the mutual information.

#### Algorithm 2: Algorithm for the local permutation method for surrogate generation.

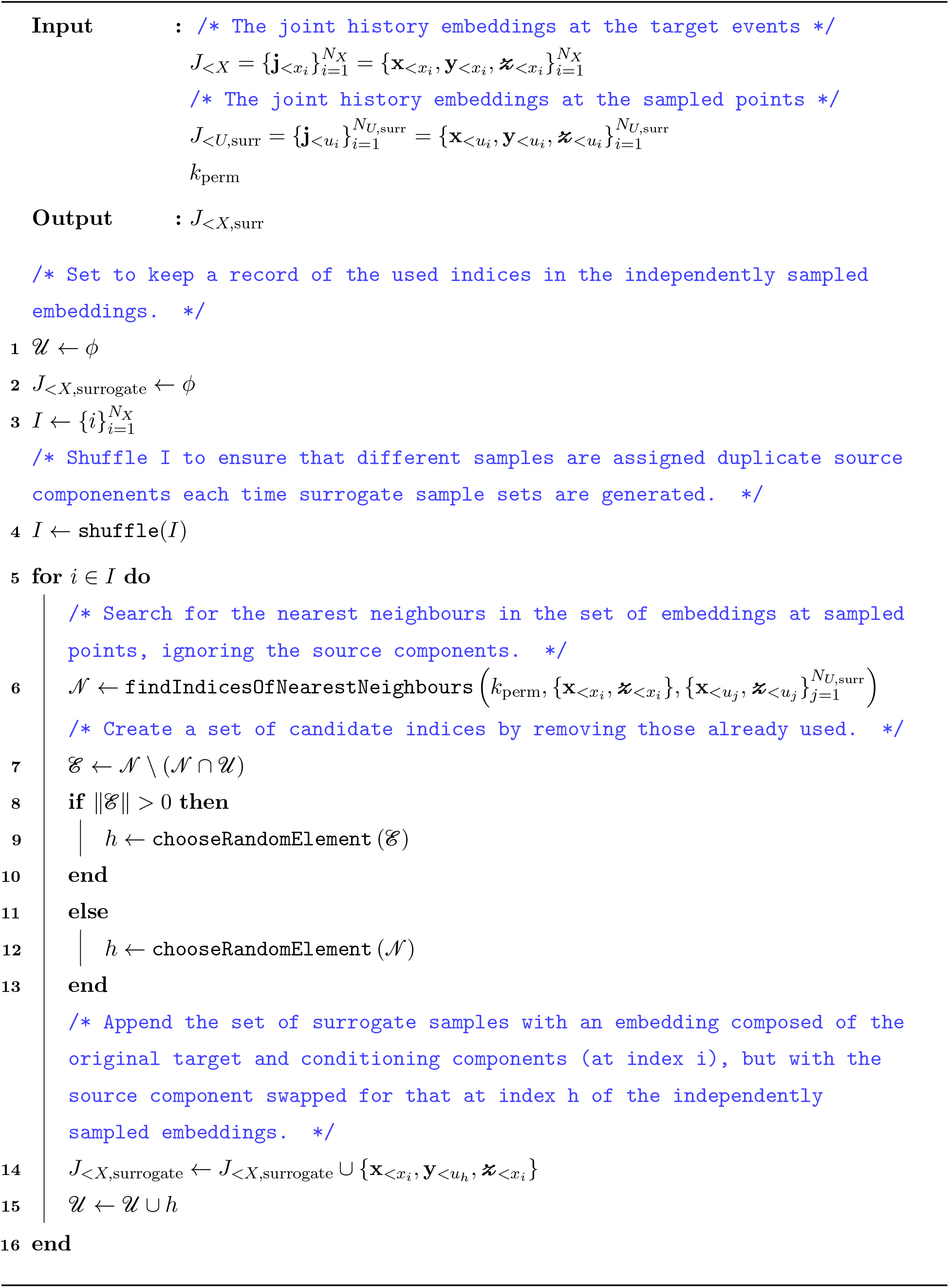

As transfer entropy is a conditional mutual information 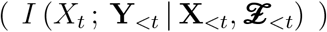, we are testing against the null hypothesis that the current state of the target *X_t_* is conditionally independent of the history of the source 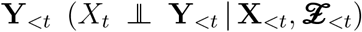. That is, the null hypothesis states that the joint distribution factorises as: 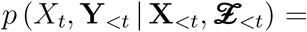 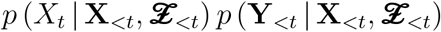.

Historically, the generation of surrogates for TE has been done by either shuffling source history embeddings or by shifting the source time series (see discussions in e.g. [4, 76]). These approaches lead to various problems. These problems stem from the fact that they destroy any relationship between the source history (**Y**_<*t*_) and both the target (**X**_<*t*_) and conditioning 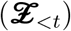 histories. As such, they are testing against the null hypothesis that the joint distribution factorises as: 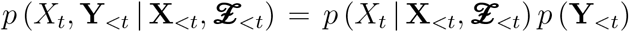. [8] The problems associated with this factorisation become particularly pronounced when we are considering a system whereby the conditioning processes 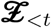 drive both the current state of the target *X*_*t*_ as well as the history of the source **Y**_<*t*_. This can lead to **Y**_<*t*_ being highly correlated with *X*_*t*_, but conditionally independent. This is the classic case of a “spurious correlation” between **Y**_<*t*_ and *X*_*t*_ being mediated through the “confounding variable” 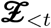. If, in such a case, we use time shifted or shuffled source surrogates to test for the significance of the TE, we will be comparing the TE measured when *X*_*t*_ and **Y**_<*t*_ are highly correlated (albeit potentially conditionally independent) with surrogates where they are independent. This subtle difference in the formulation of the null may result in a high false positive rate in a test for conditional independence. An analysis of such a system is presented in section II C. Alternately, if we can generate surrogates where the joint probability distribution factorises correctly and the relationship between **Y**_<*t*_ and the histories **X**_<*t*_ and 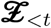 is maintained, then **Y**_<*t*_ will maintain much of its correlation with *X_t_* through the mediating variable 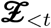. We would anticipate conditional independence tests using surrogates generated under this properly formed null to have a false positive rate closer to what we expect.

Generating surrogates for testing for conditional dependence is relatively straightforward in the case of discrete-valued conditioning variables. If we are testing for dependence between *X* and *Y* given *Z*, then, for each unique value of *Z*, we can shuffle the associated values of *Y*. This maintains the distributions *p*(*X*|*Z*) and *p*(*Y* |*Z*) whilst, for any given value of *Z*, the relationship between the associated *X* and *Y* values is destroyed.

The problem is more challenging when *Z* can take on continuous values. However, recent work by Runge [8], demonstrated the efficacy of a local permutation technique. In this approach, to generate one surrogate sample set, we separately generate a surrogate sample 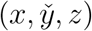 for each sample (*x, y, z*) in the original set. We find the *k*_perm_ nearest neighbours of *z* in *Z*: one of these neighbours, *z*′, is chosen at random, and *y* is swapped with the associated *y*′ to produce the surrogate sample (*x, y*′*, z*). In order to reduce the occurrence of duplicate *y* values, a set 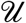 of used indices is maintained. After finding the *k*_perm_ nearest neighbours, those that have already been used are removed from the candidate set. If this results in an empty candidate set, one of the original *k*_perm_ candidates are chosen at random. Otherwise, this choice is made from the reduced set. As before then, a surrogate conditional mutual information is estimated for every surrogate sample set, and a population of such surrogate estimates provides the null distribution.

This approach needs to be adapted slightly in order to be applied to our particular case, because we have implictly removed the target variable (whether or not the target is spiking) from our samples via the novel Bayesian inversion. We can rewrite (7) as:

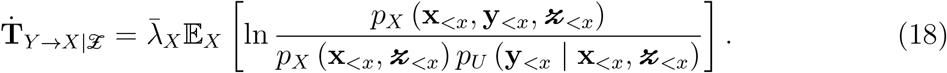

This makes it clear that we are testing whether the following factorisation holds: 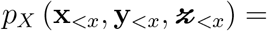 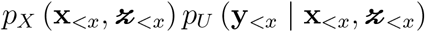 (recall the difference between probability densities at target events *p_X_* and those not conditioned at target events *p_U_*). In order to create surrogates *J*_<*X*,surr_ that conform to this null distribution, we resample a new set from our original data in a way that maintains the relationship between the source histories and the histories of the target and conditioning processes, but decouples (only) the source histories from target events. (As above, simply shuffling the source histories across *J*_<*X*_ or shifting the source events does not properly maintain the relationship of the source to the target and conditioning histories). The procedure to achieve this is detailed in algorithm 2. We start with the samples at target events 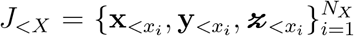 and resample the source components 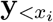 as follows. We first construct a new set 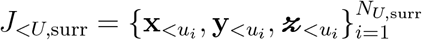 from the set *U*_surr_ of *N*_*U*,surr_ points sampled independently of events in the target. This set is constructed in the same manner as *J_<U_*, although we might choose to change the number of sample points (*N*_*U*,surr_ ≠ *N_U_*) at which the embeddings are constructed. For each original sample 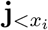 from *J*_<*X*_ then, we then find the nearest neighbours 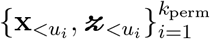 of 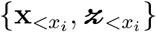 in *J*_<*U*,surr_ (line 6 of algorithm 2), select 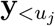 randomly from amongst the *k*_perm_ nearest neighbours (line 9 or 12), and add a sample 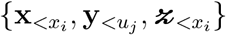 to *J*_<*X*,surr_ (line 14). The construction of such a sample is also displayed in Fig. 10. Similar to Runge [8], we also keep a record of used indices in order to reduce the incidence of duplicate 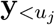 (line 15). For each redrawn surrogate sample set *J*_<*X*,surr_ a surrogate conditional mutual information is estimated (utilising the same *J*_<*U*_ selected independently of the target events as was used for the original TE estimate) following the algorithm outlined earlier; the population of such surrogate estimates provides the null distribution as before.

**FIG. 10:**
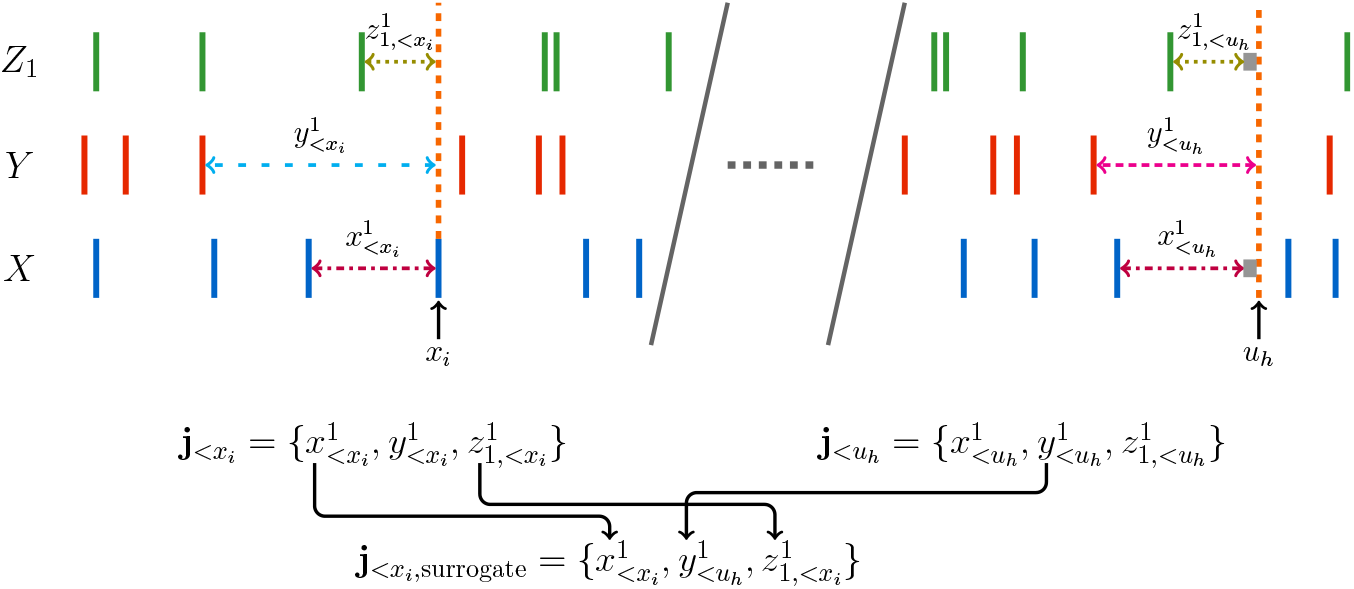
Diagrammatic representation of the local permutation surrogate generation scheme. For our chosen sample 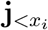 we find a 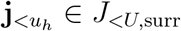 where we have that the 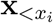 component of 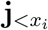 is similar to the 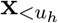 component of 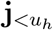 and 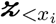 component of 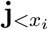 is similar to the 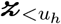 component of 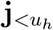. We then form a single surrogate sample by combining the 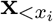 and 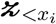 components of 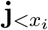 with the 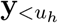 component of 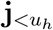. Corresponding colours of lines indicates corresponding length. The grey boxes indicate a small delta.

Finally, note an additional subtlety for dynamic correlation exclusion for the surrogate calculations. Samples in the surrogate calculations will have had their history components originating from two different time windows. One will be from the construction of the original sample and the other from the sample with which the source component was swapped. A record is kept of both these exclusion windows and, during neighbour searches, points are excluded if their exclusion windows intersect either of the exclusion windows of the surrogate history embedding.

### C. Implementation

The algorithms 2 and 1 as well as all experiments were implemented in the Julia language. All code will be made available upon publication. Implementations of *k*-NN information-theoretic estimators have commonly made use of KD-trees to speed up the nearest neigh-bour searches [76]. A popular Julia nearest neighbours library (NearestNeighbors.jl [77]) was modified such that checks for dynamic exclusion windows (see section IV A 6) were performed during the KD-tree searches when considering adding neighbours to the candidate set. The implementation of the algorithms is available at the following repository: github.com/dpshorten/CoTETE.jl. Scripts to run the experiments in the paper can be found here: github.com/dpshorten/CoTETEexperiments.

## V. ACKNOWLEDGEMENTS

We would like to thank Mike Li for performing preliminary benchmarking of the performance of the discrete-time estimator on point processes.

## VI. AUTHOR CONTRIBUTIONS

**Conceptualization:** David P. Shorten, Richard E. Spinney and Joseph T. Lizier

**Funding acquisition:** Joseph T. Lizier

**Methodology:** David P. Shorten, Richard E. Spinney and Joseph T. Lizier

**Software:** David P. Shorten

**Supervision:** Richard E. Spinney and Joseph T. Lizier

**Writing - original draft:** David P. Shorten

**Writing - review & editing:** David P. Shorten, Richard E. Spinney and Joseph T. Lizier

## VII. FUNDING STATEMENT

JL was supported through the Australian Research Council DECRA Fellowship grant DE160100630 and through The University of Sydney Research Accelerator (SOAR) prize program. High performance computing facilities provided by The University of Sydney (artemis) have contributed to the research results reported within this paper.

## Appendix A Simulation of the Pyloric Circuit of the STG

The simulation approach used follows very closely that presented in [61, 62, 78]. The only significant deviation is the addition of a noise term.

We modelled the neurons of the of the pyloric circuit using a conductance-based model.

The membrane potential (*V*) evolves according to

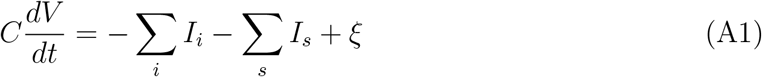

*C* = 0.628 nF is the membrane conductance. *ξ* is a noise term. Each current is specified by

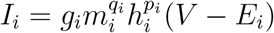

*E_i_* is the reversal potential and its values are listed in table I. The reversal potentials associated with the calcium channels are not listed as these are calculated according to the Nernst equation. Specifically, 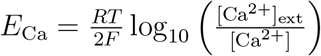 where *R* = 8.314 J K^−1^ mol^−1^ is the universal gas constant, *R* = 293.3 K is the and *F* = 96 485.332 12 C mol^−1^ is Faraday’s constant. [Ca^2+^]_ext_ = 3 mm is the extracellular Ca^2+^ concentration and [Ca^2+^] is the intracellular Ca^2+^ concentration. The intracellular Ca^2+^ concentration evolves according to

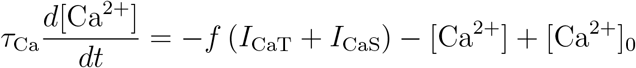

**TABLE I.** Parameters and functions used in the conductance based model.

| Current | E | $p$ | $q$ | $m_\infty$ | $h_\infty$ |
| --- | --- | --- | --- | --- | --- |
| $I_{Na}$ | 50 | 3 | 1 | $\frac{1}{1 + \exp\left(\frac{V+25.5}{-5.29}\right)}$ | $\frac{1}{1 + \exp\left(\frac{V+48.9}{5.18}\right)}$ |
| $I_{CaT}$ | | 3 | 1 | $\frac{1}{1 + \exp\left(\frac{V+27.1}{-7.2}\right)}$ | $\frac{1}{1 + \exp\left(\frac{V+32.1}{5.5}\right)}$ |
| $I_{CaS}$ | | 3 | 1 | $\frac{1}{1 + \exp\left(\frac{V+33}{-8.1}\right)}$ | $\frac{1}{1 + \exp\left(\frac{V+60}{6.2}\right)}$ |
| $I_A$ | -80 | 3 | 1 | $\frac{1}{1 + \exp\left(\frac{V+27.2}{-8.7}\right)}$ | $\frac{1}{1 + \exp\left(\frac{V+56.9}{4.9}\right)}$ |
| $I_{KCa}$ | -80 | 4 | 0 | $\frac{[Ca]}{[Ca] + 3} \frac{1}{1 + \exp\left(\frac{V+28.3}{-12.6}\right)}$ | |
| $I_{Kd}$ | -80 | 4 | 0 | $\frac{1}{1 + \exp\left(\frac{V+12.3}{-11.8}\right)}$ | |
| $I_H$ | -20 | 1 | 0 | $\frac{1}{1 + \exp\left(\frac{V+75}{5.5}\right)}$ | |

[Ca^2+^]_0_ = 0.05 μm is the steady-state Ca^2+^ concentration, *f* = 14.96 μm nA^−1^ and 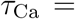 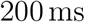

The values of *q_i_* and *p_i_* are listed in table I. The activation variables *m_i_* evolve according to

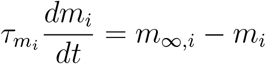

The inactivation variables *h_i_* evolve according to

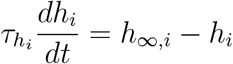

*m*_∞_,*_i_* and *h*_∞_,*_i_* are given in table I and 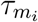 and 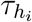 are given in table II.

**TABLE II.** Parameters and functions used in the conductance based model.

| Current | $\tau_m$ | $\tau_h$ |
| --- | --- | --- |
| $I_{Na}$ | $2.64 - \frac{2.52}{1 + \exp\left(\frac{V+120}{-25}\right)}$ | $\frac{1.34}{1 + \exp\left(\frac{V+62.9}{-10}\right)} \left[ 1.5 - \frac{42.6}{1 + \exp\left(\frac{V+34.9}{3.6}\right)} \right]$ |
| $I_{CaT}$ | $43.4 - \frac{42.6}{1 + \exp\left(\frac{V+68.1}{-20.5}\right)}$ | $210 - \frac{179.6}{1 + \exp\left(\frac{V+55}{-16.9}\right)}$ |
| $I_{CaS}$ | $2.8 + \frac{14}{\exp\left(\frac{V+27}{10}\right) + \exp\left(\frac{V+70}{-13}\right)}$ | $120 + \frac{300}{\exp\left(\frac{V+55}{9}\right) + \exp\left(\frac{V+65}{-16}\right)}$ |
| $I_A$ | $23.2 - \frac{20.8}{1 + \exp\left(\frac{V+32.9}{-15.2}\right)}$ | $77.2 - \frac{58.4}{1 + \exp\left(\frac{V+38.9}{-26.5}\right)}$ |
| $I_{KCa}$ | | $180.6 - \frac{150.2}{1 + \exp\left(\frac{V+46}{-22.7}\right)}$ |
| $I_{Kd}$ | | $14.4 - \frac{12.8}{1 + \exp\left(\frac{V+28.3}{-19.2}\right)}$ |
| $I_H$ | | $\frac{2}{\exp\left(\frac{V+169.7}{-11.6}\right) + \exp\left(\frac{V-26.7}{14.3}\right)}$ |

The synaptic currents are specified by

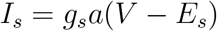

*E_s_* is the synaptic reversal potential. It was set to −70 mV for glutamatergic synapses and −80 mV for cholinergic synapses. The activation variable *a_s_* evolves according to

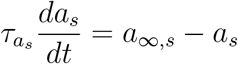

where

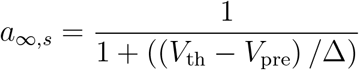

and

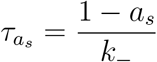

Δ = 5 mV provides the slope of the activation curve. *V*_th_ = −35 mV is the half activation potential of the synapse. *V*_pre_ is the membrane potential of the presynaptic neuron. *k*_−_ is the rate constant for the transmitter-receptor dissociation rate. For the glutamatergic synapses we used *k*_−_ = 0.025 ms and for the cholinergic synapses we used *k*_−_ = 0.01 ms.

Simulations of the pyloric rhythm can be run with fixed maximum conductance values *g_i_* and *g_s_*, as in [61]. However, it was found that these models were less robust to the addition of a noise term. It was, therefore, decided to use adaptive conductances as described in [62]. Each conductance evolved according to:

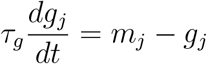

where

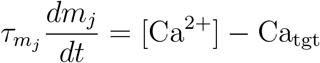

*τ_g_* = 100 ms was common across all channels. The time constants 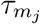 are listed in table III.. The time constants provided in [62] were not used as these produce a pyloric rhythm with unrealistically short period. Instead, the approach presented in [62] for arriving at conductance time constants from desired conductance values was used. The conductance values in [61] were used as these desired values. Specifically, we used the values presented in table II for AB/PD 1, LP 2 and PY 1. The time constant 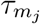 associated with *g_j_* was set as 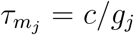. *c* is a constant with units of seconds that was adjusted by hand so that the activity converged in a reasonable amount of time and the time constants were of the same order of magnitude as those provided in [62]. As in [62], the leak conductances were fixed. They were set at 0 for the AB/PD neuron, and 0.0628 μS for the PY neuron and 0.1256 μS for the LP neuron.

**TABLE III.**
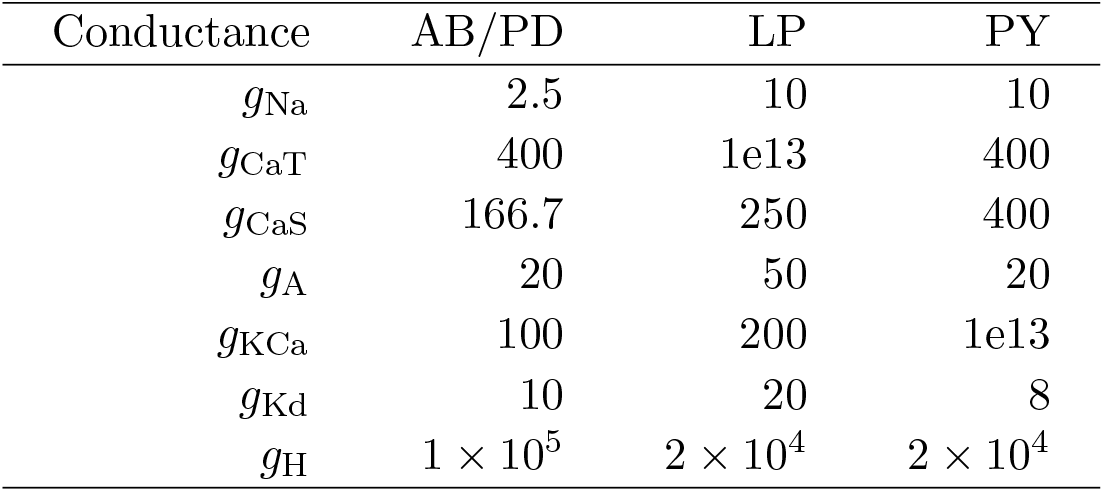
The conductance time constants 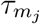. All values are in seconds.

**TABLE IV.**
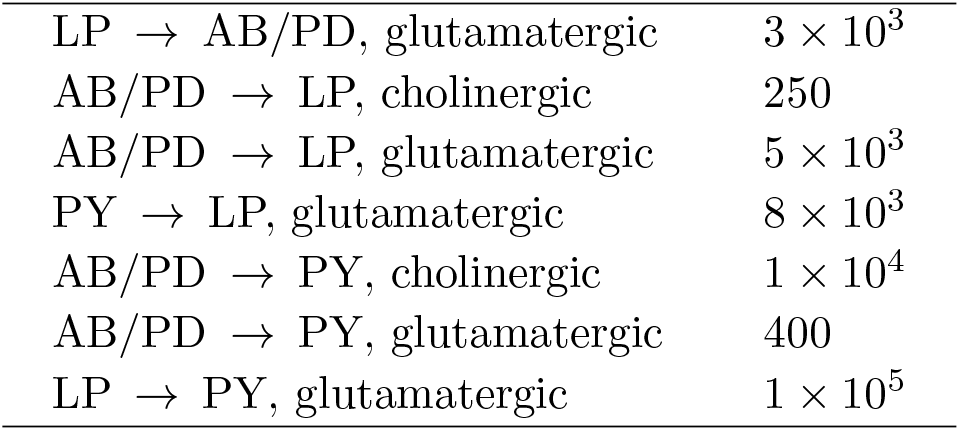
The conductance time constants 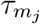 for the synapses. All values are in seconds.

|  |  |
| --- | --- |
| LP → AB/PD, glutamatergic | $3 \times 10^3$ |
| AB/PD → LP, cholinergic | 250 |
| AB/PD → LP, glutamatergic | $5 \times 10^3$ |
| PY → LP, glutamatergic | $8 \times 10^3$ |
| AB/PD → PY, cholinergic | $1 \times 10^4$ |
| AB/PD → PY, glutamatergic | 400 |
| LP → PY, glutamatergic | $1 \times 10^5$ |

The TE approach to network inference will not work in a fully deterministic system. As such, noise was added to the system. There are a number of techniques for adding noise to conductance-based models [79]. The simplest such technique is to add noise to the currents in (A1). Although this is not a biophysically realistic method, it has been shown to produce resulting behaviours which closely match those produced by more realistic techniques [79, 80]. As such, we decided to make use of this procedure in our simulations. The associated noise term is shown in (A1). The noise was generated using an AR(1) process

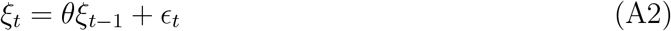

We used the parameter value of *θ* = 0.01. *ϵ_t_* was distributed normally with mean 0 and standard deviation 8 × 10^−13^

A simulation timestep of Δ*t* = 0.01 ms was used.

